# A statistical framework for differential pseudotime analysis with multiple single-cell RNA-seq samples

**DOI:** 10.1101/2021.07.10.451910

**Authors:** Wenpin Hou, Zhicheng Ji, Zeyu Chen, E. John Wherry, Stephanie C. Hicks, Hongkai Ji

## Abstract

Pseudotime analysis with single-cell RNA-sequencing (scRNA-seq) data has been widely used to study dynamic gene regulatory programs along continuous biological processes. While many computational methods have been developed to infer the pseudo-temporal trajectories of cells within a biological sample, methods that compare pseudo-temporal patterns with multiple samples (or replicates) across different experimental conditions are lacking. Lamian is a comprehensive and statistically-rigorous computational framework for differential multi-sample pseudotime analysis. It can be used to identify changes in a biological process associated with sample covariates, such as different biological conditions, and also to detect changes in gene expression, cell density, and topology of a pseudotemporal trajectory. Unlike existing methods that ignore sample variability, Lamian draws statistical inference after accounting for cross-sample variability and hence substantially reduces sample-specific false discoveries that are not generalizable to new samples. Using both simulations and real scRNA-seq data, including an analysis of differential immune response programs between COVID-19 patients with different disease severity levels, we demonstrate the advantages of Lamian in decoding cellular gene expression programs in continuous biological processes.

## 1 Introduction

Single-cell RNA-sequencing (scRNA-seq) enables dissection of complex cellular programs at single-cell resolution in biological samples with heterogeneous cell compositions. When the cells in a sample come from a continuous biological process, such as a temporal or spatial process, computationally placing cells along a pseudotemporal trajectory based on their progressively changing transcriptomes is a powerful approach to reconstructing the dynamic gene expression programs of the underlying biological process. This approach, also known as *pseudotime analysis*^1–3^, is now widely used to study cell differentiation^4–6^, immune responses^7, 8^, disease development^9–12^ and many other biological systems with temporal or spatial dynamics. A systematic review and comparison of these methods can be found in a recent benchmark study^3^, but the majority of the methods were designed to infer gene expression changes along the reconstructed trajectory within one biological sample. However, scRNA-seq experiments today standardly generate data with multiple biological samples across multiple conditions. For example, a number of COVID-19 studies generated scRNA-seq data from multiple patients with differential disease severity levels^13–19^. Therefore, there is an increasing demand for methods that can simultaneously (i) take into account sample-to-sample variation and (ii) identify changes in pseudotemporal trajectories across conditions. To meet this demand, several challenges need to be addressed.

First, changes in pseudotemporal trajectories across conditions can occur in multiple ways, including (i) topological differences, such as a cell lineage along differentiation is lost (or added) in one sample group compared to another group, (ii) changes in the proportion (or density or abundance) of cells along a cell lineage across conditions, and (iii) changes in the gene expression itself along pseudotime across conditions. An ideal solution would address all three types of changes in one comprehensive statistical framework. While there currently exist pseudotime analysis methods to detect changes in gene expression along pseudotime (e.g. `Monocle`^20–22^, `TSCAN`^23^, Slingshot^24^), in cell abundance along pseudotime (e.g. `milo`^25^, `DAseq`^26^), and in trajectory lineages (e.g. `tradeSeq`^27^), none of these methods investigate changes across conditions.

A second important challenge is to incorporate sample-to-sample variation in the framework described above. Almost all existing methods ignore this variation by either only analyzing cells from a single sample or treating cells from multiple samples as if they were from a single sample. It is important to consider the variation across samples, because ignoring sample-to-sample variation can lead to identifying false discoveries that are not generalizable or replicable in independent studies with new samples.

To the best of our knowledge, there currently does not exist a comprehensive framework that provides an integrative solution to identify the three types of changes in pseudotemporal trajectories (topology, cell density, and gene expression) across experimental conditions with multiple samples per condition. To address this gap, we introduce a comprehensive and integrative statistical framework, referred to as *Lamian*, for differential multi-sample pseudotime analysis. Given scRNA-seq data from multiple biological samples with known covariates, such as age, sex, sample type, disease status, Lamian can be used to (1) construct pseudotemporal trajectories and evaluate the uncertainty of the topologies, (2) evaluate differential changes in the topological structure associated with sample covariates, (3) describe how gene expression and cell density change along the pseudotime, and (4) characterize how sample covariates modifies the pseudotemporal dynamics of gene expression and cell density. Importantly, when identifying gene expression or cell density changes, Lamian accounts for variability across biological samples. As a result, Lamian is able to more appropriately control the false discovery rate (FDR)^28^ when analyzing multi-sample data, a property not offered by other existing methods.

## 2 Results

### 2.1 Lamian: a statistical framework for differential pseudotemporal trajectory analysis in multiple samples

Lamian consists of four modules tackling different aspects of multi-sample pseudotime analysis (Fig. 1). The input for Lamian includes (1) low-dimensional space representation, such as principal components analysis (PCA) or uniform manifold approximation and projection (UMAP)^29^, of the scRNA-seq data from multiple samples that have been harmonized and embedded into a common space using methods such as Seurat^30^ or Harmony^31^, (2) the normalized scRNA-seq gene expression matrices, and (3) sample-level metadata, such as covariate information corresponding to samples’ characteristics or biological groups information. Advantages of Lamain compared to existing methods (Table S1) include evaluating tree topology uncertainty and differential topology, and identifying gene expression and cell density changes associated with sample covariates while accounting for sample-level variability.

**Figure 1.**
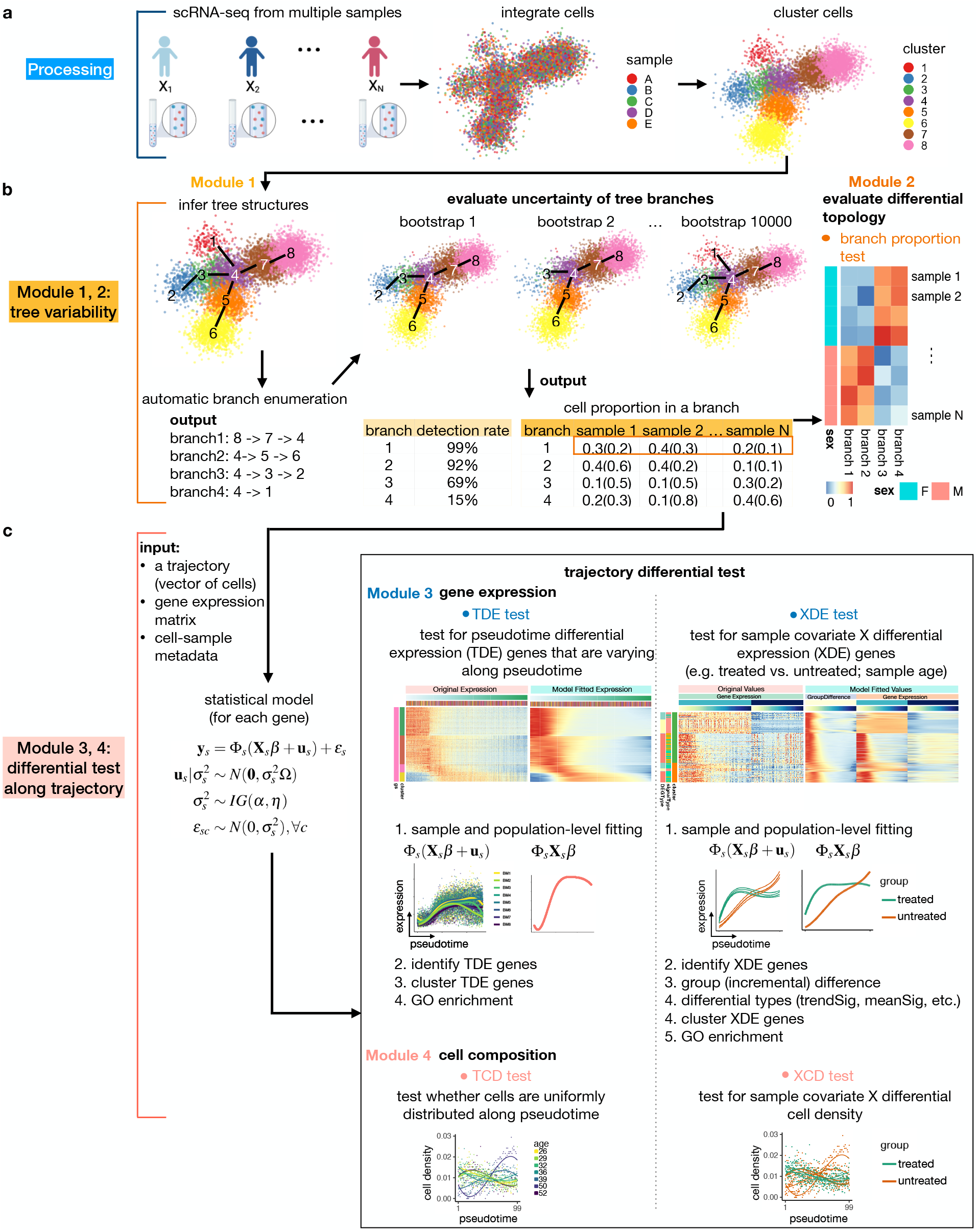
Overview of Lamain: a statistical framework for differential pseudotemporal trajectory analysis with multiple samples. **(a)** Using integrated and harmonized scRNA-seq data across multiple samples, Lamian first group cells into clusters. **(b)** Clustered data is used to infer a pseudotemporal trajectory followed by automatically enumerating all pseudotemporal paths and branches. The uncertainty of tree branches are quantified using a *detection rate* in bootstrap resampling framework (Module 1), followed by quantifying the variability of branches across samples and identifying differences in branching structure across conditions (Module 2). **(c)** For each tree branch or pseudotemporal path, Lamian can identify two types of differential expression (DE): DE along pseudotime (TDE) and DE associated with a sample covariate (XDE) (Module 3). Similarly, Lamian can also identify changes in cell density, both along pseudotime (TCD) and associated with a sample covariate (XCD) (Module 4).

#### 2.1.1 Constructing a pseudotemporal trajectory and quantifying the uncertainty of the tree branches

Module 1 of Lamian uses the harmonized data to construct a pseudotemporal trajectory and then quantifies the uncertainty of tree branches using bootstrap resampling. First, cells from all samples are jointly clustered (Fig. 1a), and the cluster-based minimum spanning tree (cMST) approach described in TSCAN^23^ is used to construct a pseudotemporal trajectory. The tree can have multiple branches, allowing one to model multiple lineages of a dynamic process. Next, after users specify a tree node as the start of pseudotime or marker genes that should highly express at the start of pseudotime, Lamian will automatically enumerate all pseudotemporal paths and branches. Then, it evaluates the uncertainty of each branch by quantifying a metric we refer to as the *detection rate*, which is defined as the probability that a tree branch can be detected in repeated bootstrap samplings of cells (Fig. 1b).

#### 2.1.2 Differential tree topology across conditions

Module 2 of Lamian first identifies variation in tree topology across samples and then assesses if there are differential topological changes associated with sample covariates (Fig. 1b). For each sample, Lamian calculates the proportion of cells in each tree branch, referred to as *branch cell proportion*. Because a zero or low proportion can reflect absence or depletion of a branch, changes in tree topology can be described using branch cell proportion changes. With multiple samples, Lamian characterizes the cross-sample variation of each branch by estimating the variance of the branch cell proportion across samples. Furthermore, regression models can be fit to test whether the branch cell proportion is associated with sample covariates. This allows one to identify tree topology changes between different conditions, for example in a case-control cohort.

#### 2.1.3 Differential gene expression analysis along a pseudotemporal trajectory

Given a path or branch along a pseudotemporal trajectory, the scRNA-seq gene expression matrices from multiple samples, and sample-level covariate information, Module 3 of Lamian identifies differentially expressed (DE) genes (Fig. 1c). There are two types of DE tests. First, the *TDE test* evaluates whether a gene’s activity as a function of pseudotime *t*, denoted as *f* (*t*), is a constant (*H*_0_ : *f* (*t*) = *c*), with the goal to identify genes whose activities change along pseudotime (*H*_1_ : *f* (*t*) ≠ *c*). Here, TDE refers to *pseudotime differential expression*. In contrast, the *XDE test* evaluates for each gene whether the pseudotemporal activity *f* (*t*) is associated with a sample-level covariate, such as whether *f* (*t*) is different between healthy and disease samples. Here, XDE refers to *covariate X differential expression*. Currently, existing pseudotime methods, such as Monocle, Slingshot and TSCAN only detect TDE, but not XDE. Lamian is the first integrative framework to provide both TDE and XDE for multiple sample analyses. For each XDE gene, Lamian further evaluates whether the sample covariate shifts the mean of *f* (*t*) (referred to as a *mean shift*) or changes the functional form of *f* (*t*) (referred to as a *trend difference*) or both. Additionally, unsupervised *k*-means clustering is applied to DE genes to group and summarize major differential gene patterns. In all DE tests, Lamian accounts for sample-to-sample variation directly into the estimation framework, whereas the other methods do not. Consequently, Lamian is able to control the false discovery rate (FDR)^28^ compared to existing methods that ignore sample-to-sample variation which leads to identifying false discoveries that are not generalizable in new samples.

#### 2.1.4 Differential cell density analysis along a pseudotemporal trajectory

Similar to gene expression, Module 4 of Lamian tests whether cells’ density along pseudotime is uniformly distributed or not (*TCD test*), and if it is associated with a sample covariate (*XCD test*). This may be used to study dynamic processes, such as cell expansion in immune response or how disease changes the pseudotemporal cell density pattern.

### 2.2 Lamian estimates tree topology stability and accurately detects differential tree topology

We first illustrate Modules 1 and 2 of Lamian using a Human Cell Atlas (HCA)^32, 33^ 10x Genomics scRNA-seq dataset consisting of bone marrow samples, referred to as HCA-BM, from 8 donors and a total of 32,819 cells.

#### 2.2.1 Estimate tree topology stability using HCA-BM data

First, we assess the tree topology stability (Module 1). Bone marrow contains hematopoietic stem cells (HSCs) differentiating into different blood cell types. Applying TSCAN to the harmonized bone marrow data, we identified 6 cell clusters (Fig. 2a), which form a minimum spanning tree with three branches, corresponding to the three major lineages of HSC differentiation - myeloid, erythroid, and lymphoid (Fig. 2b). We confirmed these lineages with marker genes (Fig. 2c). Specifically, HSCs are mostly in cluster 5, as indicated by high CD34 expression (Fig. 2c). By setting cluster 5 as the origin, we obtained three pseudotemporal paths (Fig. 2a: the path of cluster 5 → 1; 5 → 6 → 2; 5 → 3 → 4). Lamian uses repeated bootstrap sampling of cells along the branches to calculate a detection rate. In the HCA-BM data, these three branches can be detected in 93.8% (5 → 1), 95.3% (5 → 6 → 2) and 61.5% (5 → 3 → 4) in all bootstrap samples (or with a detection rate = 0.938, 0.953 and 0.615), suggesting that they are real branches rather than random noise.

**Figure 2.**
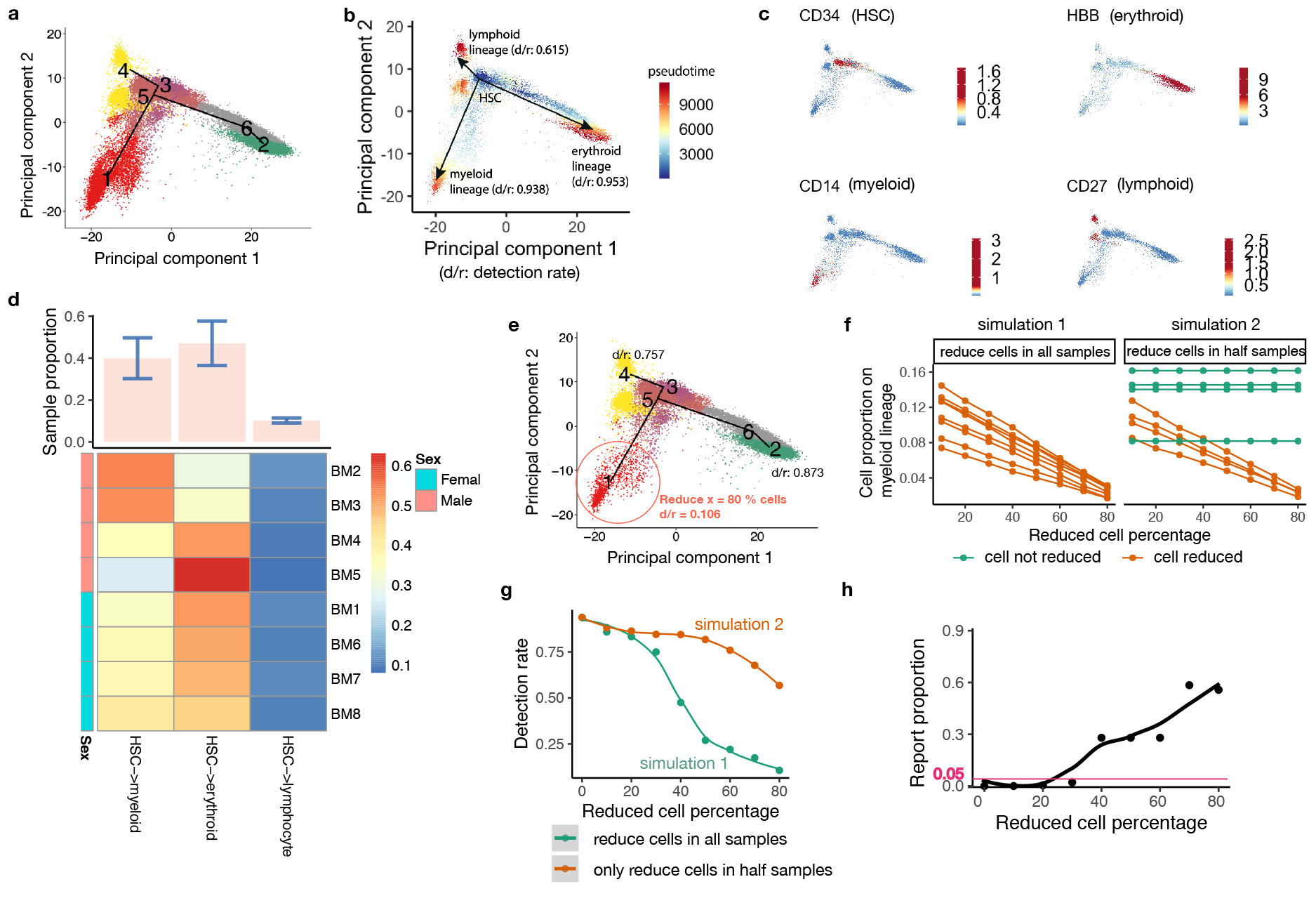
Lamain estimates tree topology stability (Module 1) and tests differential tree topology between sexes (Module 2) in the HCA bone marrow data^32, 33^. **(a)** Inferred tree topology using eight integrated scRNA-seq bone marrow samples displayed in the first two principal components (PCs). Each dot is a cell colored by the cluster label (*k* = 6). **(b)** Similar to (a), but cells are colored by the inferred pseudotime. The estimated detection rates (d/r) are shown for three tree branches which correspond to three major lineages of hematopoietic stem cell (HSC) differentiation. **(c)** Similar to (a), but cells are colored by the expression of lineage-specific marker genes. **(d)** Heatmap of sample-level branch cell proportion (the number of cells in each branch divided by the total number of cells in a sample). The barplot on the top shows the mean and standard deviation (SD) of the branch cell proportion across samples for each lineage. Samples are also colored by sex. **(e)** Tree topology and detection rates after randomly removing *x* = 80% cells on the myeloid lineage (branch 5 → 1) as an illustration of simulation. **(f)** Simulations are conducted in two ways by either removing certain percentage of cells (x-axis) along the myeloid lineage across all eight samples (simulation 1: left) or removing cells in only half of the samples (simulation 2: right). **(g)** Lamian-reported detection rate of the myeloid lineage (y-axis) after removing different percentage of cells (x-axis) in the two simulations. **(h)** In simulation 2, the difference between two sample groups increases as the reduced cell percentage *x* increases. For each *x*, 10,000 simulations were run. The proportion of p-values smaller than the significance cutoff 0.05 in the simulations (y-axis) is shown as a function of the reduced cell percentage (x-axis).

#### 2.2.2 Differential tree topology tests using HCA-BM data

Next, we assess the variability in the branch cell proportions across samples and between conditions (Module 2). Using all 8 donors, the branch cell proportion is 41.1%, 48.4% and 10.5% for the myeloid, erythroid and lymphoid branches, respectively. Of note, the proportions show variation across donors (proportion Mean (SD) = 0.41 (0.10) for myeloid, 0.48 (0.11) for erythroid, 0.11 (0.01) for lymphoid). Lamain uses a two-sided *t*-test to assess if there is a statistically significant difference in the tree topology (i.e. branch cell proportion) between two sample groups. However in the HCA-BM data, comparing the branch cell proportion between male and female donors did not show significant differences along the myeloid, erythroid, and lympoid lineages (*p*-values = 0.53, 0.60, 0.60, respectively), suggesting that there is no significant change in tree topology between the two sexes (Fig. 2d).

#### 2.2.3 Lamian accurately characterizes stability and differences in tree topology

To demonstrate the validity of Lamian’s topology stability and differential topology analysis, we performed two sets of simulations. In Simulation 1, we reduced the number of cells in the myeloid lineage in the HCA-BM data by subsampling (Fig. 2e,f). As expected, decreasing the number of cells decreased the detection rate for the myeloid branch (Fig. 2g). For example, when 80% cells were reduced, the detection rate dropped to 0.106 (Fig. 2e,g). Hence, the detection rate provides a reasonable measure for quantifying the certainty (or uncertainty) conveyed by the data about the presence of a branch.

In Simulation 2, we reduced the number of cells in the myeloid lineage in four out of the eight samples (Fig. 2f). As the number of cells decreased, the detection rate of the myeloid branch again decreased, but at a much slower rate compared to Simulation 1 (Fig. 2g). We found that conditional on the branch being detected, our differential topology test (Module 2) was able to detect differences in the branch cell proportion between the two groups of samples in this simulation scenario. Most importantly, it controls the probability of false positives (type I error rate) when there are no differences (i.e. removing no cells or 0% of cells) and also has increasing statistical power to detect true positives as we increase the percent of cells removed in half of the samples (Fig. 2h).

### 2.3 Lamian comprehensively detects differential pseudotemporal gene expression and cell density

We next illustrate how Lamian adjusts for sample-to-sample variation to identify differential gene expression (Module 3: TDE and XDE tests) and differential cell density (Module 4: TCD and XCD tests) along pseudotime using the eight samples in the HCA-BM dataset.

#### 2.3.1 Detect differentially expressed genes along pseudotime using HCA-BM data

First, we ask which genes are varying along pseudotime (Module 3: TDE test). Applying the TDE test with a 5% FDR cutoff, Lamian identified 8,475, 7,454 and 8,953 TDE genes for the myeloid, erythroid and lymphoid lineage, respectively (Fig.3a-c). Among the TDEs, we found known lineage markers corresponding to each lineage, such as CD14 for myeloid, HBB for erythroid, and CD3D, CD19, CD27 for lymphoid. Hence, TDE genes can be used to identify branch lineages in the tree topology. Unsupervised clustering of TDE genes and gene ontology (GO) analysis revealed the cascade of the transcriptional program (Fig. 3a-c, Fig. S1a-f). For example, as HSCs differentiate to the erythroid lineage, the TDE genes with initially high expression but low expression at the end are enriched in CD8-positive, alpha-beta T cell activation, whereas genes with increasing expression along pseudotime are enriched in oxygen transport (Fig. S1c,d). Meanwhile, for the lymphoid lineage, the TDE genes with high expression initially but low expression at the end are enriched in platelet degranulation, whereas genes with increasing expression along pseudotime are enriched in T cell differentiation (Fig. S1e,f).

**Figure 3.**
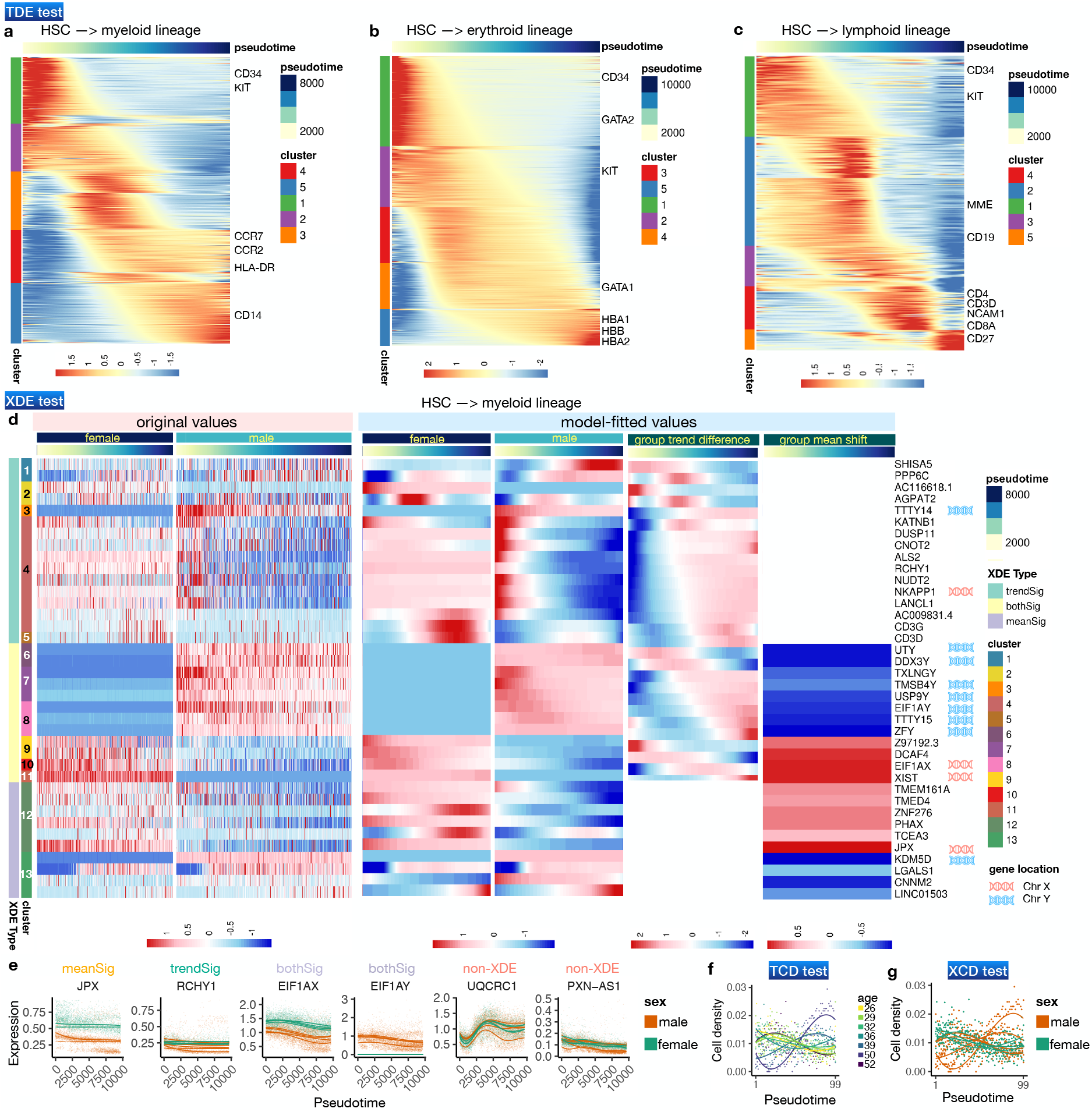
Lamian supports comprehensive analysis of differential gene expression (Module 3: TDE and XDE tests) and cell density (Module 4: TCD and XCD tests) along pseudotime in the HCA bone marrow data. **(a-c)** Heatmap of model-fitted expression values for TDE genes (rows, FDR < 0.05) along HSC differentiating to myeloid (a), erythroid (b), and lymphoid (c) inferred pseudotime lineages. Rows are clustered using *k*-means clustering and ordered by differential gene patterns. **(d)** Heatmaps of Lamian-detected differentially expressed (XDE) genes along the myeloid lineage (rows, FDR < 0.05) between male and female (Module 3). The heatmaps only show the 38/43 XDE genes annotated as having either significant mean shift or trend difference (i.e., 10 *meanSig*, 16 *trendSig*, and 12 *bothSig* genes). The other XDE genes (*otherSig* genes: neither *FDR_trend_* < 0.05 nor *FDR_mean_* < 0.05) are not included in the heatmaps. The six heatmaps from the left to right correspond to raw normalized gene expression along pseudotime for each sex (left 2 heatmaps), model fitted gene expression along pseudotime in each sex (middle 2 heatmaps), trend difference and mean shift between female and male along pseudotime (right 2 heatmaps). Genes from chromosomes X and Y are labeled. **(e)** Example XDE (and non-XDE) gene expression along the myeloid lineage with significant mean shift (*meanSig*), trend difference (*trendSig*), and both (*bothSig*). The fitted curve for each sample is also shown. **(f)** The model-fitted cell density pseudotemporal patterns in myeloid lineage in TCD test. Each curve depicts the model-fitted values for one sample. **(g)** Similar to (f) but curves are colored by sex used in the XCD test. The corresponding figures for (d-g) in the erythroid and lymphoid lineages can be found in Fig. S2 (Module 3: XDE tests) and Fig. S3 (Module 4: TCD and XCD tests).

#### 2.3.2 Detect differentially expressed genes associated with a covariate along pseudotime using HCA-BM data

Next, we tested whether there are differential gene expression patterns along pseudotime associated with sex as a covariate (Module 3: XDE test). For each gene, Lamian reports three FDRs: (1) *FDR_overall_* corresponds to testing if a gene is XDE (*overall test*), (2) *FDR_trend_* corresponds to testing if a XDE gene has significant trend difference associated with the sample covariate (*trend test*), and (3) *FDR_mean_* corresponds to testing if a XDE gene has significant mean shift associated with the covariate (*mean test*). In addition, there are two other categories: both mean and trend differences (*bothSig*), or neither mean or trend differences (*otherSig*). Using the XDE test, Lamian identified 43, 32 and 29 genes (*overall test*) with significant differences (at the 5% *FDR_overall_* cutoff) between male and female along the myeloid (Fig. 3d), erythroid (Fig. S2a), and lymphoid (Fig. S2b) lineages, respectively. Next, Lamian further annotated the XDE genes into the gene patterns described above. For the myeloid lineage, this results in 10 genes with mean shift only, 16 genes with trend difference only, and 12 genes with significant changes both in mean and trend (Fig. 3d,e). Among the XDE genes, 33% (*N*=14) are from chromosome X and Y, representing a significant enrichment in sex chromosomes (Fig. 3d, permutation test *p*-value = 0.0036 for chromosome X and *p* < 10^−5^ for chromosome Y, see Methods). Notably, among the genes that show significant mean shift (with or without trend difference), 12 genes have higher mean expression in males and they consist of 8 genes on Y chromosome and 4 genes on autosomes. Likewise, 10 genes have higher mean in females and they consist of 3 genes on X chromosome and 7 genes on autosomes (Fig. 3d). Unsupervised clustering of XDE genes revealed cascades of their dynamic transcriptional programs. For example, among genes with trend difference only, the difference in SHISA5 expression between female and male was positive at the beginning and negative at the end of the pseudotime, whereas the difference in DUSP11 was negative at the beginning and positive at the end (Fig. 3d). Analyses of the erythroid and lymphoid lineages yielded similar results (Fig. S2a,b).

#### 2.3.3 Detect changes in cell density along pseudotime using HCA-BM data

Finally, we tested for changes in cell density both along the pseudotime (Module 4: TCD test) and whether these patterns were associated with sex as a sample covariate (Module 4: XCD test). The TCD test shows that cell density changed significantly along all three lineages (myeloid: Fig. 3f; erythroid: Fig. S3a; and lymphoid: Fig. S3c) (all *p*-values after adjusting for multiple testing are < 10^−200^), although in this dataset it is unclear whether the cell density change was due to technical sampling bias (e.g. certain cell types are easier to sample) or real biology. In the XCD test, we did not find significant differences in cell density along pseudotime between male and female (myeloid: Fig. 3g; erythroid: Fig. S3b; and lymphoid: Fig. S3d).

### 2.4 Lamian is more powerful than existing methods to detect differences while controlling the FDR by accounting for sample-level variation

In this section, we use simulated and real scRNA-seq data to demonstrate how Lamian is more powerful than existing methods to detect gene expression differences that are associated with a covariate (Module 3: XDE test). We also demonstrate how incorporating the the sample-to-sample variation into the differential gene expression test along psuedotime (Module 3: TDE test) leads to less false discoveries compared to existing methods that also perform TDE detection.

#### 2.4.1 Lamian controls the FDR in differential gene expression tests associated with sample-level covariates

First, we created a null data set based on the HCA-BM data (described in detail in Methods)). Briefly, we first randomly partitioned the eight HCA-BM samples into two groups and removed the group differences to create a dataset where we do not expect any XDE genes between the two groups (Fig. 4a). When Lamian was applied to detect group differences, no XDE genes were reported. Building upon the null data set above, we then introduced *in silico* spike-in differential signals with varying strengths and pseudotemporal patterns between the two sample groups to a random set of genes (details in Methods). In this way, we know which genes are XDE genes and whether they have mean shift, trend difference, or both (Fig. 4b).

**Figure 4.**
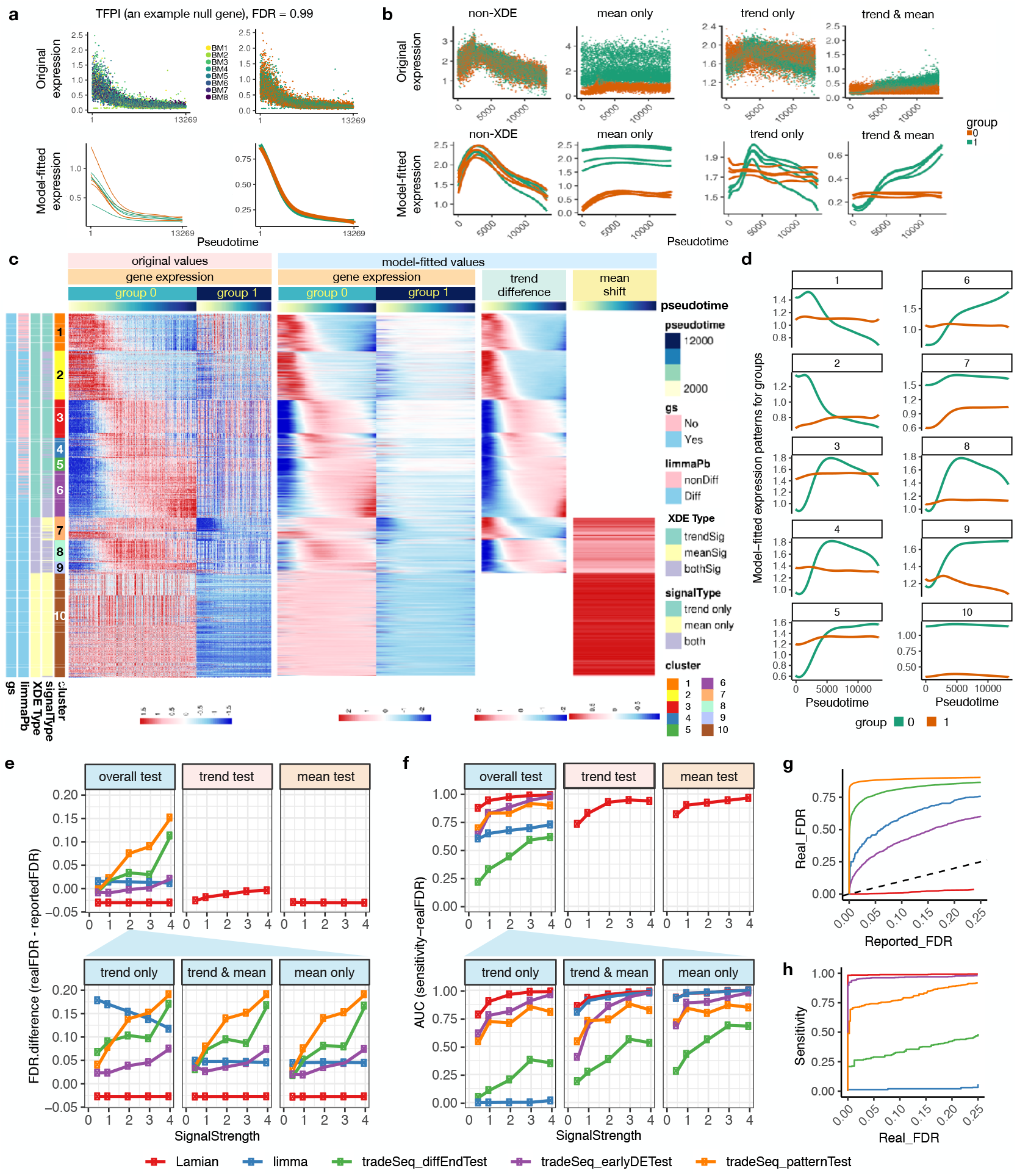
Evaluation of the FDR control and statistical power for detecting differential genes associated with sample covariate (XDE). **(a)** An example null gene where no differential signals are expected along the trajectory between two groups of samples. In the top panel, each dot is a cell colored by the sample (left) or the sample group (right). The plots show the gene’s expression along pseudotime in each cell. In the bottom panel, curves are model-fitted gene expression patterns for each sample (left) or sample group (right). **(b)** Examples of non-differential (non-XDE) genes and differential (XDE) genes with only mean difference (mean only), trend difference (trend only), and both mean and trend difference (trend & mean) between two sample groups in the simulation data. For each gene, the top panel shows the expression along pseudotime in each cell. Each dot is a cell colored by the sample group. The bottom panel shows sample-level model-fitted gene expression curves along pseudotime. **(c)** Heatmaps in four white-bar-seperated panels to show the expression patterns of XDE genes (rows) by cells (columns) ordered by pseudotime. The 1st and 2nd panels show original values and model-fitted values of gene expression. Cells from the samples in group 0 and 1 are separated. The 3rd and 4th panels show the standardized model-fitted group difference (trend difference) and the mean shift between groups, where white space denotes no significant difference. **(d)** Model-fitted temporal patterns of group 1 and 0 averaged across XDE genes in each gene cluster. **(e-f)** Performance evaluation of all methods in five spike-in signal strengths settings (x-axis; signal strength increases from 0 to 4). **(e)** Plots in the first row shows the difference between the true and reported FDR for overall XDE test, trend test, and mean test. Here, the FDR difference is the difference between the area under the realFDR-reportedFDR curve and the diagonal line as illustrated in (g). Plots in the second row compare the FDR difference from different methods when gold-standard genes are stratified into trend, mean, and trend & mean differences. **(f)** Similar to (e) but compares the power using the area under sensitivity-realFDR curve as illustrated in **(h)**

Next, we applied Lamian to identify XDE genes and clustered genes based on their differential patterns (Fig. 4c-d). We compared Lamian with limma and tradeSeq, two widely used tools to detect differences in gene expression. As limma is designed to detect differential mean gene expression, we pooled all cells on a pseudotemporal path or branch to create a pseudobulk expression profile (i.e. the average expression across cells for a gene) for each sample. In this way, limma uses the pseudobulk data to detect mean differences between two sample groups. In contrast, tradeSeq (which is used by Slingshot) is a method originally developed for comparing different branches of a pseudotemporal trajectory within a single sample. Here, we tailored the function to compare the same branch in a pseudotemporal trajectory between two samples. Since tradeSeq does not consider cross-sample variability, cells from replicate samples are pooled and treated as if they come from a single sample.

For all three tests (*overall test*, *trend test*, *mean test*), and across all signal strength levels, the real FDR was smaller than the FDR reported by Lamian, demonstrating that Lamian was able to conservatively estimate FDR (Fig. 4e,g). Limma and tradeSeq do not report separate FDRs for mean and trend differences. Limma reports an overall FDR for each gene. TradeSeq can be run to detect different types of DE: earlyDETest identifies genes that show expression difference in early pseudotime; patternTest identifies genes that show expression difference along all pseudotime that are equally-spaced; diffEndTest compare the average expression at the end stage of pseudotime. It assigns an FDR for each test. Unlike Lamian, both limma and tradeSeq underestimated the real FDR: the difference between the real FDR and their reported FDR was positive in most cases (Fig. 4e). We also stratified XDE genes into three groups - mean shift only, trend difference only, and both mean and trend differences - based on their true states. Within each stratum, the *FDR_overall_* reported by Lamian conservatively estimated the real FDR, whereas limma and tradeSeq underestimated the real FDR (Fig. 4e).

#### 2.4.2 Lamian is powerful to detect differences in gene expression associated with sample-level covariates

We further compared the statistical power of different methods via the sensitivity-real FDR curve and the area under the curve (AUC) (Fig. 4f, h). The power of detecting XDE genes by Lamian increased with increasing signal strength, both for detecting XDE genes overall or for detecting a specific class of XDE genes (Fig. 4f). For detecting all XDE genes (*overall test*), both limma and tradeSeq had lower power compared to Lamian (Fig. 4f). When XDE genes are stratified, limma had comparable power to Lamian for detecting XDE genes with mean shift (i.e. mean shift only or both mean and trend differences) but had zero power to detect genes with trend difference only. TradeSeq had lower power in all XDE gene categories (Fig. 4f).

In addition to our simulation studies, we compared the output from Lamian, limma and tradeSeq using the real HCA-BM dataset to detect sex differences (Fig. 5). For the myeloid lineage, limma detected 5 XDE genes and all of them were found by Lamian. Lamian reported an additional 38 genes not found by limma (25 with trend difference, 9 with mean shift only) (Fig. 5a). TradeSeq, on the other hand reported 3,677 XDE genes. However, a closer examination of the results from tradeSeq indicates that at least a subset of these genes are false positives (Fig. 5b). For example, both BCLAF1 and CHPT1 were reported as XDE by tradeSeq. For each gene, when cells from replicate samples were treated as if they were from one sample, the fitted gene expression curve along pseudotime are different between male and female, which explains why tradeSeq reported the gene as XDE. However, when the gene expression curve is fitted within each sample, the variation among replicate samples is much bigger than the difference between male and female and hence there is no real statistically significant sex difference (Fig. 5b). As a comparison, Fig. 5c illustrates example genes reported by Lamian. Here, the sex difference is clear even after accounting for sample variability. Indeed, while XDE genes reported by Lamian and limma both significantly overlap with both chromosome X and chromosome Y, XDE genes reported by tradeSeq did not show a statistically significant association with chromosome X (Fig. 5c). Note that XDE genes found by Lamian but not limma also significantly overlap with sex chromosomes, suggesting that these genes are indeed sex related (Fig. 5d). The performance of Lamian on the other two lineages was similar (Fig. S4). Collectively, our analyses demonstrate that Lamian provides the unique ability to detect XDE genes not offered by other methods.

**Figure 5.**
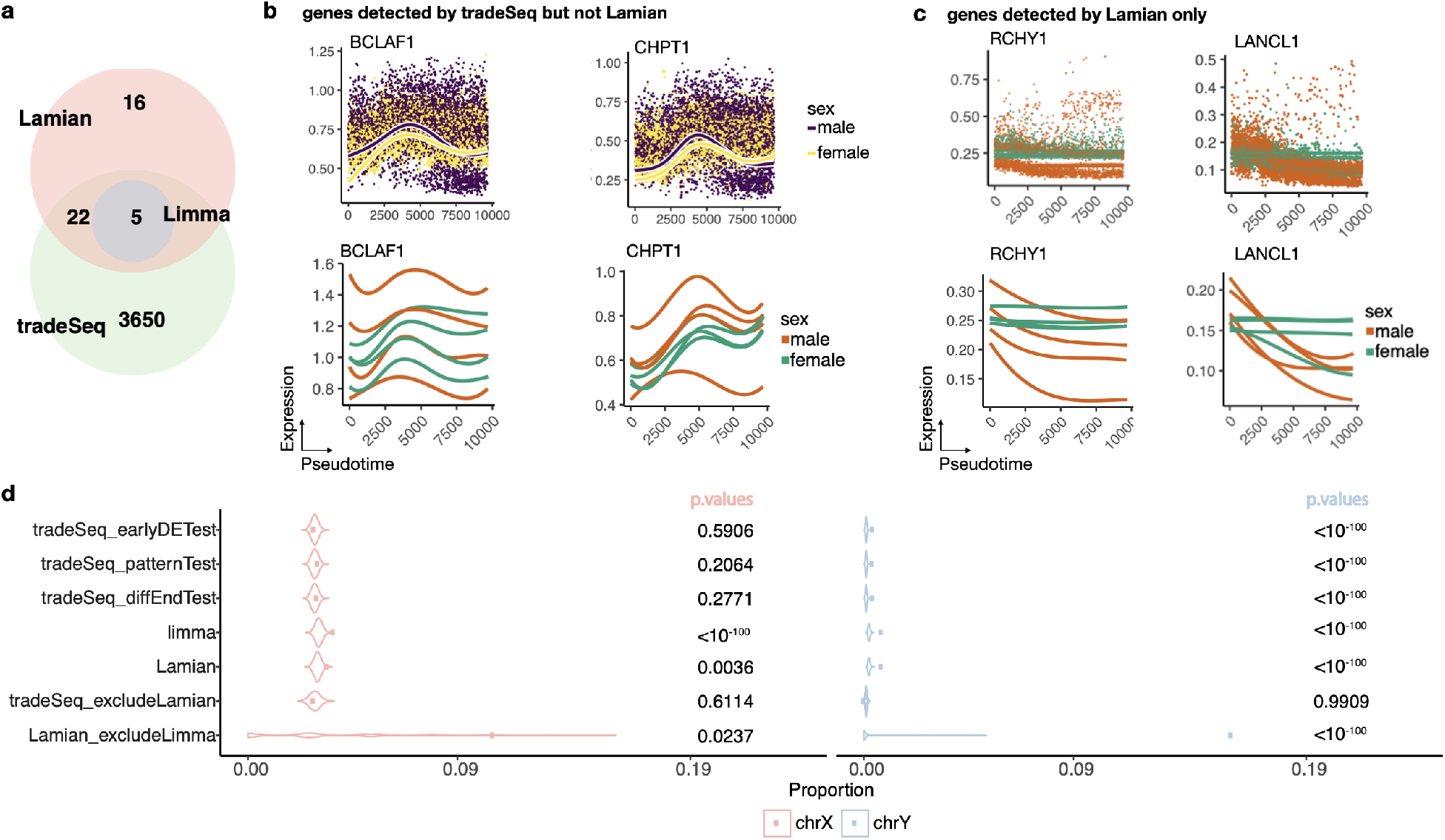
Comparison of sex-associated XDE detection of Lamian and other methods in HCA bone marrow data. **(a)** A Venn diagram of the number of XDE genes detected by Lamian, limma and tradeSeq (pooled three types of tests) in the myeloid lineage when comparing female and male samples. **(b)** Two example genes detected by tradeSeq but not Lamian. Purple-yellow plots display the patterns output by tradeSeq while the green-brown plots display the pseudotemporal patterns for each sample (curve) fitted by Lamian. **(c)** Two example genes only detected by Lamian. The first row shows gene expression in each cell along pseudotime. Each dot is a cell. The second row shows gene expression curve fitted by Lamian in each sample. **(d)** Overlap between XDE genes reported by different methods and sex chromosome genes as a gold standard (left: chromosome X; right: chromosome Y). The overlap was calculated for top *N* XDE genes with different *N*s. The mean of the overlap across all *N*s was used as the observed overlap statistic (see an illustration in Fig.S4a). Violin plots show the permutation null distribution used to determine the statistical significance of the observed overlap statistics (shown as dots), and the p-values are shown on the right of each plot.

#### 2.4.3 Incorporating sample-level variation reduces false positives in TDE detection compared to existing methods

In addition to detecting differentially expressed genes along pseudotime that are associated with a covariate, Lamian can also detect differentially expressed genes along pseudotime without any covariate information (Module 3: TDE test). In this case, there are existing methods, such as Monocle, Slingshot, tradeSeq and TSCAN that perform a similar test. However, we note that, compared to existing methods, Lamian is unique in that it incorporates sample-to-sample variability into the statistical estimation framework. Using simulated data with multiple samples, we found that Lamian, compared to existing methods, controls the FDR, while also maintaining strong statistical power for TDE detection (Supplementary Notes, Fig. S5).

Finally, similar to DE analysis, our evaluation also shows that Lamian can accurately detect TCD and XCD with a well-controlled type I error rate and high statistical power (Supplementary Notes, Fig. S6).

### 2.5 Lamian analysis of COVID-19 scRNA-seq data identifies differential CD8 T cell transcriptional programs during a critical stage of disease severity transition

We applied Lamian to a COVID-19 peripheral blood mononuclear cell (PBMC) 10x Genomics scRNA-seq dataset obtained from a recent study^34^. The COVID-19 disease severity of a patient may progress from mild to moderate to severe. It was reported that the mild to moderate transition is a critical stage with rapid immune landscape changes that may determine the trajectory of disease progression^34^. CD8+ T cell activation is an important component of COVID-19 patients’ immune response to the infection. By analyzing scRNA-seq data from 66 mild and 48 moderate COVID-19 patients, we examined the CD8+ T cell activation program in these patients and asked how it changes during the mild-to-moderate disease severity transition.

First, we constructed a pseudotemporal trajectory using a total of 55,953 naive and CD8+ T cells identified from the harmonized PBMC scRNA-seq data (Fig. 6a, see details in Methods). The trajectory contains only one path without branch, thus we skip evaluating the tree branch uncertainty and differential topology. Applying TDE test, Lamian identified 2,195 TDE genes which were grouped into five clusters (Fig. 6b). Examination of these genes’ dynamic expression patterns show that the inferred pseudotemporal trajectory reflects the CD8+ T cell activation process. For example, known naïve/memory T cell associated genes including TCF7, SELL and IL7R were found in cluster 1 (Fig. 6b,c). Genes in this cluster showed decreasing expression along pseudotime, consistent with the loss of quiescent characteristics over the activation process. Genes such as JUNB and CD7 are responsible in the induction of differentiation into effectors and thus catch up expression shortly in cluster 2. Genes in this cluster 2 also include early activation marker CD69, GZMK and AP-1 family members (e.g. JUNB, JUN), suggesting that this cluster plays a role in the cell fate switch from effector memory T cells to terminal effector T cell phase. By contrast, genes in clusters 4 and 5 both show increasing expression along pseudotime, with cluster 5 reaching its peak expression later than cluster 4. We found that genes encoding functional effector molecules such as CCL5 and IFNG are enriched in cluster 4, and cluster 5 is enriched in both functional activation features such as GZMB, TBX21 and CX3CR1 and terminal differentiation gene features such as GNLY, CD244 and CD38 (Fig. 6c).

**Figure 6.**
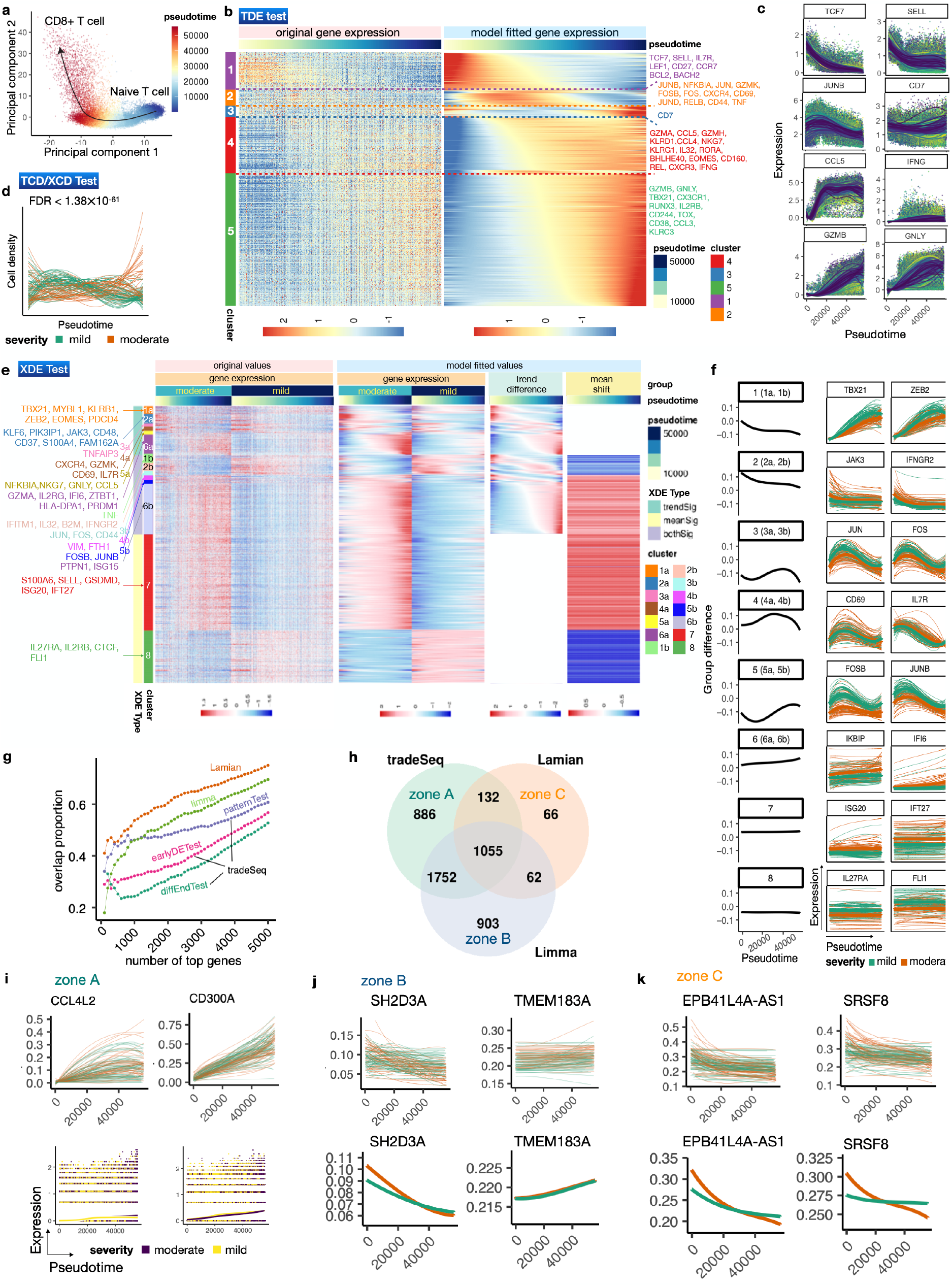
Lamian analysis of COVID-19 samples identifies differential genes related to T-cell activation and inflammation between mild and moderate patients. **(a)** PC plot of the CD8+ T cells colored by its pseudotime reflecting the T cell activation process. **(b)** As the output of TDE test, this heatmap shows the original gene expression (left) and model-fitted gene expression (right) of TDE genes along the pseudotime from naive to activated T cells as in (a). **(c)** Example TDE genes from different TDE gene clusters shown in (b). Each dot is a cell. Each curve is a sample’s pseudo-temporal pattern fitted by Lamian. **(d)** Pseudo-temporal pattern of the cell density for each sample represented by a curve colored by the patient’s severity level. FDR reported by XCD test is also shown which indicates significant difference between mild and moderate samples. **(e)** Heatmaps showing the pseudotemporal expression patterns of XDE genes (rows). Cells (columns) from moderate and mild patients are plotted separately and ordered by the pseudotime. White bars are used to separate original temporal gene expression in each group, model-fitted temporal gene expression, group trend difference (moderate group minus mild group), and group mean shift. Example genes related to immune and inflammation process are marked on the left-hand side of the heatmap. **(f)** Each XDE gene cluster in (e) is shown with the averaged group difference (left: black) and two example genes (right: green-mild, brown-moderate). For each example gene, the thin and thick curves represent its pseudo-temporal pattern for each sample and for the whole sample group, respectively. **(g)** Samples are randomly partitioned into two datasets. For each method, the proportion of top *N* XDE genes that overlap between the two datasets (*y*-axis) is shown as a function of *N* (*x*-axis). **(h)** A Venn diagram showing the number of XDE genes reported by each method. **(i-k)** Example genes that are reported only by tradeSeq (i), limma (j), or Lamian (k). The first row in each plot displays sample-level patterns. The second row of (i) shows the patterns fitted by tradeSeq, and those in (j-k) show the group patterns obtained by Lamian.

We next investigated differences in the CD8+ T cell activation program between mild and moderate patients. The analysis of cell density using XCD test shows that the abundance of activated effector T cells is significantly increased in moderate compared to mild disease (FDR = 1.38 × 10^−61^, Fig. 6d). The analysis of gene expression using XDE test identifies 1,315 XDE genes, which were grouped into 14 clusters (Fig. 6e). The first 12 clusters contain genes with pseudo-temporal trend differences (including bothSig and trendSig), and their trend differences follow 6 major patterns (Fig. 6f, e.g. cluster 2a and 2b have the same trend difference pattern, but cluster 2a has no significant mean shift whereas cluster 2b has significant mean shift). The last 2 clusters contain genes with mean shift only. In cluster 1, TBET (encoded by TBX21) and ZEB2 are major transcription factors (TF) for CD8 T cell effector responses^35–38^ and drive IFNG production. Genes in this cluster tend to have lower expression in moderate patients compared to mild patients and the magnitude of difference increases along the pseudotime (Fig. 6e,f), suggesting that mild patients have a more robust functional effector CD8 T cell response. In cluster 6 (incl. 6a and 6b), several interferon stimulated genes such as IFI6 and ISG15 as well as terminal differentiation transcription factor BLIMP-1(encoded by PRDM1)^39^ become increasingly more upregulated in moderate patients compared to mild patients along pseudotime, suggesting that a stronger inflammation in moderate patients drives CD8 T cell termination. Together, these data indicate that compared to mild disease, CD8 T cells in moderate COVID-19 patients are programmed to be less functional effector-like and more terminally differentiated. This is consistent with previous observation that comparing to the COVID-19-recovered donors, ongoing disease patients show a more TEMRA differentiation with less T-bet+ functional effector CD8 T cells^40^.

We further compared Lamian with limma and tradeSeq for detecting XDE genes. We first randomly partitioned the COVID samples into two sets and detected XDE genes between mild and moderate samples within each set. We then examined the proportion of overlap between the two XDE gene lists. By applying Lamian, we achieved the highest overlap proportion between the two partitioned data sets. Among the remaining methods, the patternTest in tradeSeq performed slightly better than the other tests in tradeSeq and limma in the top 1000 genes, but limma catched up after top 1000 genes and ranked the second best after Lamian (Fig. 6g). This suggests that XDE genes identified by Lamian are most reproducible when analyzing different sets of samples.

Using all samples in the COVID-19 dataset, tradeSeq reported 3,825 XDE genes including 886 that were solely detected by tradeSeq (Fig.6h). A closer examination of the sample-level pseudotemporal curves shows that XDE genes detected only by tradeSeq consist of a large number of false positives where the mild and moderate samples were indeed mixed together due to large sample-level variation (Fig. 6i). Limma reported 3,772 genes, including 903 that were solely detected by limma. The sample-level curves show that many of the genes reported only by limma did not show clear group differences (Fig. 6j). Lamian reported 1,315 XDE genes, including 66 that were solely detected by Lamian. For these genes, group differences cannot be explained away by the sample-level variation (Fig. 6h,j).

Collectively, our analyses demonstrate that Lamian provides a powerful tool for identifying differences associated with covariates that the other methods do not offer. The COVID analysis also demonstrates the value of using multi-sample differential pseudotime analysis for understanding dynamic gene expression programs in a disease.

## 3 Discussion

Our results demonstrate that Lamian provides a systematic solution to multi-sample pseudotime analysis capable of detecting topology, gene expression and cell density differences between different conditions. Lamian evaluates statistical significance after accounting for cross-sample variability which is important to filter out false discoveries that are not generalizable to new samples. Lamian is a free and open source R package with a modular structure. While we demonstrated its default pipeline in this article, users can replace certain modules by their own data or algorithm. For example, instead of constructing pseudotime using TSCAN in Lamian, they can provide their own pseudotemporal trajectories constructed using other algorithms and directly start analysis from module 2.

Lamian is computationally tractable. For analyzing the HCA bone marrow dataset with 32,819 cells and 8 samples, it took 4.1 hours to run the whole pipeline (0.1 h for trajectory variability, 1.5h for XDE detection and 1.5h for TDE detection, 0.01h for cell density test) on a computer cluster with 25 CPUs (2.5 GHz CPU and at most 163 GB RAM). For analyzing 39,512 CD8 T cells in the COVID dataset with 114 samples, it took 37 hours to run the whole analysis pipeline.

Currently, the statistical model in Lamian is formulated for scRNA-seq data. However, its general principle and statistical framework may be applicable to other data types such as single-cell ATAC-seq data as well, although the other data types may have different data characteristics that requires one to tailor the model accordingly. These extensions will be a topic for future research.

## 4 Method

### 4.1 Data

#### Human Cell Atlas bone marrow dataset (HCA-BM)

The raw count matrix of bone marrow scRNA-seq data sequenced in 10x Genomics platform from 8 healthy donors were downloaded from the Human Cell Atlas (HCA) data portal^32, 33^. The raw data consist of 42,925 genes and 290,861 cells. Cells with fewer than 5,000 reads, fewer than 1,000 expressed genes (i.e. genes with nonzero read count), or more than 10% of reads mapped to the mitochondrial genome were deemed as low quality and filtered out. We also filtered out genes that were expressed in less than 0.1% of all cells. This results in a data matrix of 22,401 genes × 32,819 cells used for subsequent analyses.

#### COVID19 dataset (COVID-Su)

The raw count matrices of 256 PBMC 10x Genomics scRNA-seq samples from 139 COVID-19 patients were downloaded from E-MTAB-9357 (https://www.ebi.ac.uk/arrayexpress/experiments/E-MTAB-9357/)^34^. We filtered out cells with fewer than 2,000 reads or 500 expressed genes or more than 10% mitochondrial reads. We also filtered out samples with fewer than 500 cells. `Seurat`(v.3.2.1)^30^ was applied to process, integrate data across samples and perform the cellular clustering with default settings. Cell types were annotated based on known marker genes. CD8+ T cells were identified using CD3D expression > 1 log-scaled library-size-normalized `SAVER`-imputed read counts and CD8A expression > 1 criterion. Samples with fewer than 100 CD8+ T cells were filtered out. Among the total of 161 samples that passed the filters, we focused on analyzing samples from 66 mild and 48 moderate patients subsequently.

### 4.2 Data preprocessing

In each dataset, `Seurat`(v.3.2.1)^30^ was used to integrate multiple samples. For differential expression (DE) analysis, `SAVER` was used to impute gene expression values to address the drop-outs in the data. All DE methods used imputed values except `tradeSeq` since it requires count values as inputs. Principal Component Analysis (PCA) and Uniform Manifold Approximation and Projection (UMAP)^29^ were used for visualization, and they were both run using default settings.

### 4.3 Constructing pseudotemporal trajectory and evaluating its uncertainty

After samples are integrated, the harmonized data are used to construct pseudotemporal trajectory using a cluster-based minimum spanning tree (cMST) approach. *K*-means clustering is applied to cluster cells based on the top principal components (PCs) of log2-transformed library-size-normalized gene expression profiles. Trajectories are then inferred as in TSCAN by constructing a minimum spanning tree that treats cluster centers as nodes. The number of PCs and the cell cluster number are both determined using an elbow method as described in TSCAN^23^. The origin of the pseudotime is specified by users based on marker gene expression (or the origin cell types if users input the cell types annotation). For example, in the bone marrow data, the cluster with the highest expression of hematopoietic stem cell (HSC) marker CD34 was set as the origin. Once the origin of the trajectory is given, one can enumerate all paths and branches. Branches are identified based on nodes with degree > 2.

For each of the branch, we characterize its uncertainty using its detection rate in 10,000 bootstrap samples. Each bootstrap sample is created by sampling cells from the original data with replacement. Cells in the bootstrap sample are used to reconstruct pseudotemporal trajectory using the same cMST approach as in the original data. The origin of the pseudotime in a bootstrap sample is determined using the cell cluster with the smallest mean of cells’ pseudotime in the original data. We then ask whether each branch in the original data is also identified in the bootstrap sample by performing pairwise comparison of branches between the original and bootstrap data. For a pair of branches (one from original data and one from bootstrap sample), we use the Jaccard index to evaluate their overlap (i.e., what percentage of cells in these two branches are shared). If the Jaccard index exceeds a cutoff, then the branch in the original data is called detected in the bootstrap sample. To determine the cutoff, a null distribution of Jaccard index is constructed by evaluating the overlap between the cells in the branch and a randomly sampled set of cells with the cell number matching those in the branch for 1,000 times. The 0.99 quantile of this null distribution is used as the cutoff. After comparing the original trajectory with all bootstrap samples, the detection rate of a branch is defined as the proportion of bootstrap samples in which the original branch can be detected.

### 4.4 Tree variability across samples and differential topology analysis

For each sample, the proportion of cells in each branch is calculated and referred to as “branch cell proportion”. For each branch, the variance of branch cell proportion across samples is reported to characterize its cross-sample variability. To test differential topology, for each branch we fit a regression model using the branch cell proportion as the dependent variable and using the sample covariates specified by users as the independent variables. Statistical significance of the association between a sample covariate and the branch cell proportion is determined by testing whether the corresponding regression coefficient is zero using two-sided *t*-test. The *p*-values are adjusted for multiple testing using the Benjamini-Hochberg procedure to obtain false discovery rates (FDRs)^28^. By default, FDR≤ 0.05 is used as the significance cutoff.

### 4.5 Modeling gene expression along pseudotime

Given a pseudotemporal path or branch, Lamian will describe how gene expression *Y* varies along pseudotime *t* and characterize the relationship between each gene’s pseudotemporal expression pattern *Y* (*t*) and *V* sample covariates *X*_1_, … , *X_V_* (e.g. disease status, age, etc.).

Without loss of generality, below we presents the statistical model for one gene. All other genes can be analyzed in the same way. We use lowercase letters *s* and *c* to denote sample and cell , respectively, and we use capital letter *S* to denote the total number of samples. Assume that sample *s* consists of *C_s_* cells. Let *t_sc_* be the pseudotime of cell *c* in sample *s*. Given a gene, let *y_sc_* denote its expression level in cell *c* of sample *s*. Let **x**_*s*_ = (1, *x_s_*_1_, … , *x_sV_*)^*T*^ be the realized values of covariates in sample *s*. Here, we introduced an additional term *x_s_*_0_ ≡ 1 as an intercept term for the subsequent regression model.

We model each gene’s expression pattern along pseudotime as functional curves and represent the function using a total of *K* + 1 B-spline basis functions *ϕ*_0_(*t*), *ϕ*_1_(*t*), … , *ϕ_K_* (*t*). Here *K* is the number of equidistant knots used to define B-spline bases. The gene’s functional curve in sample *s* is 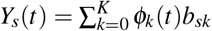. For each gene, the optimal *K* is automatically chosen by comparing values ranging from 0 to a pre-defined maximum (20 by default) and selecting the one that minimizes the Bayesian Information Criterion (BIC). The BIC for a given *K* is calculated as BIC_*K*_ = *KS* ln(∑_*s*_ *C*_*s*_) − 2∑_*s*_ *l_K,s_* + *const*. Here *const* is a constant term that does not dependent on *K* (hence irrelevant for finding optimal *K*), and *l_K,s_* is the log-likelihood of the B-spline regression for sample *s* (i.e. we fit a linear regression where the response variable is the gene expression in cells and the independent variables are the *K* + 1 *B*-spline bases).

The observed data of the gene are assumed to be generated from this unobserved function after adding cell-level random noise *ε_sc_* as follows:

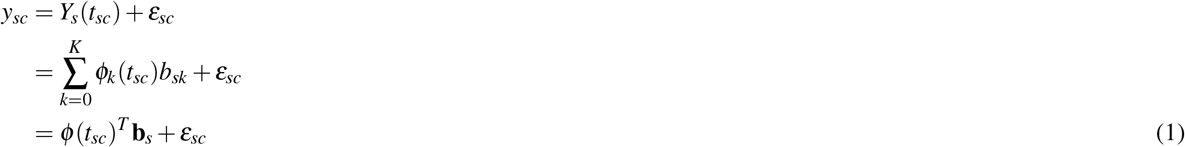

where

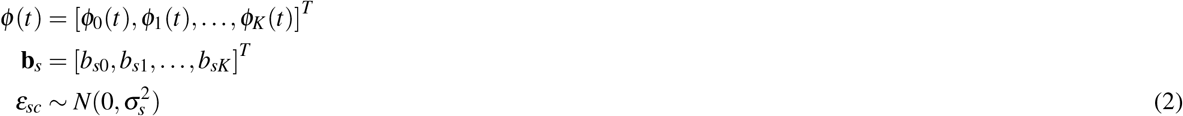

Since all samples share the same B-spline bases *ϕ* (*t*), the sample-specific temporal pattern is described via the sample-specific regression coefficients **b**_*s*_. To model the relationship between a gene’s pseudotemporal pattern *Y_s_*(*t*) and sample covariates **x**_*s*_ while accounting for sample-to-sample variability that cannot be explained by the covariates, we further assume

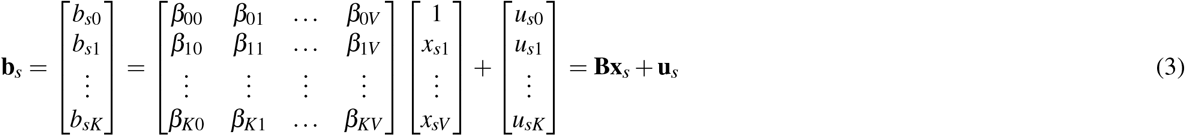

where **B** is a (*K* + 1) × (*V* + 1) matrix representing unknown fixed effects of covariates, and **u**_*s*_ is a (*K* + 1) × 1 vector representing unobserved sample-level random effects (i.e. random variations among samples with the same covariate values):

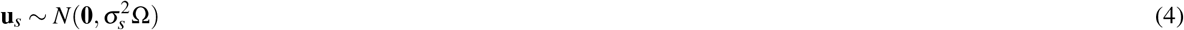

Here Ω is a (*K* + 1) × (*K* + 1) positive definite matrix. Note that the degrees of freedom for estimating sample-level covariance matrix Ω after accounting for *V* + 1 covariates are *S* – (*V* + 1) and one needs at least *K* + 1 degrees of freedom to estimate a full rank covariance matrix with dimension *K* + 1. Therefore, if the sample size *S* does not exceed *V* + *K* + 2, we do not have enough information to estimate an unconstrained Ω. In that scenario, we add a constraint by assuming Ω = *ω*^2^**I**_(*K*+1) × (*K*+1)_ where **I** represents an identity matrix. This constraint reduces the number of parameters in Ω to 1. Define

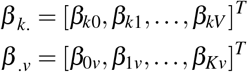

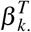. is the *k*^*th*^ row of **B**, corresponding to regression coefficients for basis *ϕ_k_*(*t*). *β _.v_* is the *v*^*th*^ column of **B**, corresponding to regression coefficients for the *v*^*th*^ covariate *X_v_*. If gene *g*’s expression pattern does not dependent on *X_v_*, then *β _.v_* = **0**.

To facilitate developing the model fitting algorithm, Equation 3 can also be rewritten in a vectorized form. Let **I** _*K*_ be a *K* × *K* identity matrix, and

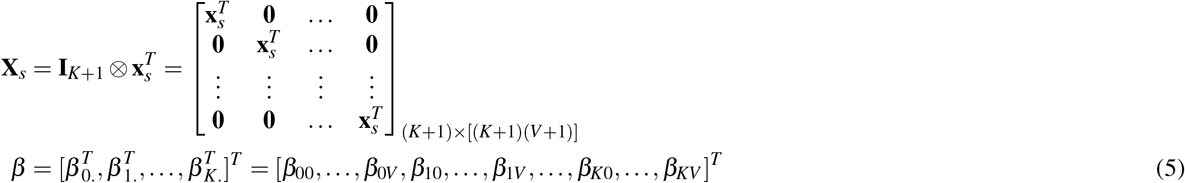

Then Equation 3 can also be written as:

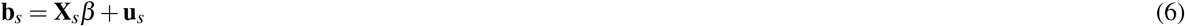

Thus, the observed data model in Equation 1 is equal to

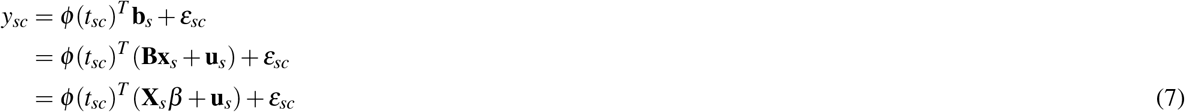

where 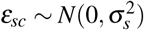 and 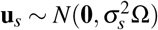. We further assume that 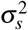 follows an inverse-Gamma distribution:

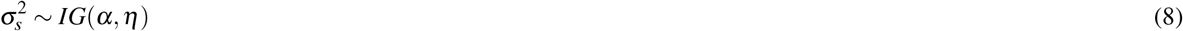

For the given gene, let **y**_*s*_ = [*y_s_*_1_, … , *y_sC_s__*]^*T*^ denote its expression in all cells in in sample *s*, *ε_s_* = [*ε_s_*_1_, … , *ε_sC_s__*]^*T*^ , and Φ_*s*_ = [*ϕ* (*t_s_*_1_), … , *ϕ* (*t_sC_s__*)]^*T*^ , then Equation 7 can also be written in a matrix form as:

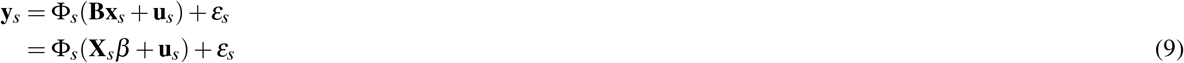

The above model can be fit using an Expectation-Maximization (EM) algorithm (see details in the **Supplementary Notes**). The algorithm can estimate the unknown parameters Θ = {*β*, Ω, *α*, *η*} and infer 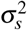 based on the observed data. Here 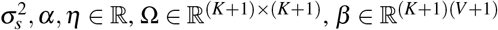.

### 4.6 Detecting differential expression associated with sample covariate (XDE)

Under the Lamian model, detecting differential expression associated with a sample covariate *X_v_* amounts to testing whether *β* _*.v*_ = [*β*_0*v*_, *β*_1*v*_, … , *β*_*Kv*_]^*T*^ = **0**. A XDE gene is a gene with *β _.v_* ≠ **0**. For a XDE gene, if *β*_0*v*_ = *β*_1*v*_ = … = *β_Kv_* = *c* (i.e. all *β_kv_*s are equal), then the effect of the covariate is to shift the gene’s pseudotemporal curve up or down by a constant *c* for every unit change in *X_v_* (because the B-spline bases satisfy 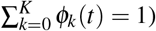. Such a gene is called XDE with mean shift only. If *β_kv_*s are not all equal for a XDE gene, then the covariate also changes the trend of the gene’s pseudotemproal curve. To systematically detect and classify XDE genes, we consider the following nested models:

- *M*_0_: *β _.v_* = [*β*_0*v*_, *β*_1*v*_, … , *β*_*Kv*_]^*T*^ = **0**.
- *M*_1_: *β* _.v_ ≠ **0** and *β*_0*v*_ = *β*_1*v*_ = … = *β_Kv_* = *c*.
- *M*_2_: *β _.v_* ≠ **0**.

We conduct the following hypothesis tests:

- *Overall XDE test*: the null model *M*_0_ is compared with the alternative model *M*_2_. Rejecting *M*_0_ implies XDE.
- *Mean test*: *M*_0_ and *M*_1_ are compared. Rejecting *M*_0_ implies mean shift.
- *Trend test*: *M*_1_ and *M*_2_ are compared. Rejecting *M*_1_ implies trend difference.

A gene is called XDE if the XDE test is significant. For a XDE gene, if the mean test is significant but the trend test is not significant, the gene is called XDE with mean shift only. If the trend test is significant but the mean test is not, then the XDE gene is called XDE with trend difference only. If both the mean test and the trend tests are significant, then the XDE gene is called XDE with both mean shift and trend difference.

To conduct a hypothesis test comparing two models, we use a permutation-based likelihood ratio test. Without loss of generality, consider comparing null model *M*_0_ versus alternative model *M*_1_ as an example (other model comparisons are handled similarly). The test statistic is the log-likelihood ratio (LLR) between *M*_1_ and *M*_0_ computed using the observed data. To construct the null distribution of the test statistics, we use a permutation approach. In each permutation, we first bootstrap the cells (keeping cell number the same as the observed data) to account for the pseudotime variability, and we then permute the values of the covariate *X_v_* among the samples. Using the permuted data, the models are refit and the LLR statistic is recomputed. Using the LLR obtained from all permutations (by default, 100 times), an empirical distribution is fitted using kernel density estimate (`base`::density()) to serve as the null distribution. The *p*-value is calculated as the tail probability of the null distribution (i.e. probability that a LLR drawn from the null distribution is equal or larger than the observed LLR). The *p*-values from all genes are adjusted for multiple testing using the Benjamini-Hochberg procedure to obtain false discovery rates (FDRs)^28^. By default, FDR*≤* 0.05 is used as the significance cutoff.

### 4.7 Detecting differential expression along pseudotime (TDE)

Unlike Lamian, most existing pseudotime methods do not detect differential expression associated with covariates (XDE). Instead, they detect differential expression along pseudotime (TDE). While our main focus is to detect XDE genes, Lamian also provides a function to detect TDE genes.

When all samples are from one group without covariate, the Equation 3 becomes

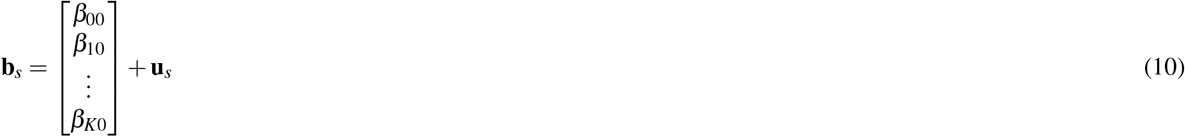

Note that 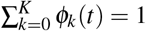. Thus, if *β*_00_ = *β*_10_ = … = *β_K_*_0_ = *c* (i.e. all *β_k_*_0_s are equal), then the pseudotemporal pattern shared by samples is *ϕ* (*t*)^*T*^ *β* _.0_ = *c*, which is a constant that does not change along pseudotime. Therefore, TDE detection can be formulated as comparing the following two models:

- *H*_0_: *β_k_*_0_ (*k* = 0, 1, … , *K*) are all equal
- *H*_1_: *β_k_*_0_ (*k* = 0, 1, … , *K*) are not necessarily all equal

This yields the following hypothesis test:

- *TDE test*: *H*_0_ and *H*_1_ are compared. Rejecting *H*_0_ implies differential expression along pseudotime (TDE).

The TDE test can also be generalized to account for sample covariates. With covariates, the compared models become:

- *H*_0_: *β_kv_* (*k* = 0, 1, … , *K*) within each column of **B** in Equation 3 are equal (i.e. *β _.v_* = *c_v_***1** where *v* = 0, 1, … , *V* and **1** represents a *K* + 1 vector with all elements equal to 1)
- *H*_1_: No constraint on **B**

The hypothesis test is conducted using a permutation-based likelihood ratio test. We first compute the log-likelihood ratio (LLR) between *H*_1_ and *H*_0_ as the test statistic using observed data. We then construct the null distribution of LLR using permutations. In each permutation, we first bootstrap the cells to to account for pseudotime variability, and we then permute the pseudotime of the cells within each sample. Using the permuted data, the models are refit and the LLR statistic is recomputed. The null distribution is derived by applying the kernal density estimate (`base`::density()) to the empirical LLR statistics obtained from all permutations (by default, for 100 times). *P*-value is calculated as the tail probability of the empirical distribution. The *p*-values from all genes are adjusted for multiple testing using the Benjamini-Hochberg procedure to obtain FDR^28^. By default, FDR*≤* 0.05 is used as the significance cutoff.

### 4.8 EM algorithm for fitting the Lamian model

The algorithm used to fit the Lamian model is provided in Supplementary Notes in detail.

### 4.9 Analysis of cell density changes

Given a pseudotemporal path or branch, we divide the pseudotime from 0 to its maximum into 100 consecutive intervals of equal lengths. The number of cells in each interval *t* and sample *s* is counted and denoted as *r_st_*. One approach to modeling cell density changes is to model *r_st_* using a count distribution (e.g. Poisson or Negative binomial) with mean *L_s_λ_st_* where *L_s_* is a sample-specific normalizing constant corresponding to the total cell number on the pseudotemporal path. One can then model log *λ_st_* as functional curves using B-spline bases similar to the gene expression model. Fitting such a model, however, requires algorithms such as Markov Chain Monte Carlo which makes this approach less appealing computationally. We therefore use an alternative and simpler approach in which *r_st_ /L_s_* is modeled in the same way as the gene expression model in equation 1 (i.e. treating time interval *t* as cell and treating *r_st_ /L_s_* in the same way as *y_sc_*). In this way, testing if the cell density changes along pseudotime (*TCD test*) or if a sample covariate changes the pseudotemporal cell density curves (*XCD test*) can be handled following the same procedure for TDE and XDE tests. This approach is more computationally efficient and yields reasonable results empirically in our benchmark data.

### 4.10 Comparisons with existing methods

#### 4.10.1 XDE detection

For detecting differential expression associated with covariates, we compared Lamian with `tradeSeq` (v.1.1.23) and `limma` (v.3.40.6). We applied tradeSeq by considering the cells belonging to two groups as those belonging to two lineages. The cell weights on each group were set as 0.99 and 0.01 respectively. We then fit the models by running the fitGAM() function with the default setting. All three types of tests for between-lineage comparisons were included. Specifically, earlyDETest(), diffEndTest() and patternTest() were applied to identify early drivers of differentiation, differentiated markers and expression patterns over pseudotime, respectively. `limma` was applied by pooling each sample as a pseudobulk. Its functions lmFit(), eBayes(), and topTable() were used to perform the test.

#### 4.10.2 TDE detection

For detecting differential expression along pseudotime, we compared Lamian with `Monocle2` (v.2.14.0), `Monocle3`^22^ (v.3.0.2.1), `tradeSeq`^27^ (v.1.1.23) and `TSCAN`^23^ (v.1.7.0). All methods other than Lamian treat cells from all samples as if they were from one sample. `Monocle2` performs the testing with an approximate *χ*^2^ likelihood ratio test. In this test, generalized additive models (GAMs) are applied to fit the gene expression against pseudotime as a full model while the null model considers the gene expression is a constant along pseudotime. `Monocle3` performs trajectory inference on the coordinates from uniform manifold approximation and projection (UMAP) and then implements the Moran’s I test to identify genes whose expression is associated with pseudotime with statistical significance. `TSCAN` applies the same fitting and testing method as `Monocle2` except that `TSCAN` uses `MGCV` package and `Monocle2` applies `VGAM` package. `tradeSeq` is used by `slingshot`^24^ to identify dynamic genes along pseudotime. Both tests designed for within-lineage comparisons in `tradeSeq` were included (startVsEndTest() and associationTest()).

#### 4.10.3 Significance cutoff

All *p*-values reported by each method were adjusted for multiple-testing using the Benjamini-Hochberg procedure to obtain false discovery rates (FDRs)^28^. By default, FDR≤ 0.05 is used as the significance cutoff.

### 4.11 Simulations

#### 4.11.1 XDE detection

We first created null simulation data where we do not expect any XDE genes. The simulation was based on the 13,269 cells on the erythroid branch in the real HCA-BM data described in Section 4.1. For the null simulation in Fig. 4a, the eight bone marrow scRNA-seq samples were randomly partitioned into two groups (group 0 and 1). Next, to remove any group differences for a given gene, we divided the pseudotime into 100 non-overlapping intervals of equal lengths. Within each interval and within each sample group, we calculated the median of the gene’s normalized expression. For cells in the sample group with lower median value, we added their expression with the difference of median expression between the two groups so that the two groups have similar expression values.

Building upon the null dataset above, we then introduced *in silico* spike-in differential signals with varying strengths and pseudotemporal patterns between the two sample groups to a random set of genes. This spike-in simulation data set was used in Fig. 4b-h. We randomly selected 20% (1814) genes as the gold standard XDE genes (gs genes) and randomly assigned them to 3 groups: trend difference only, mean shift only, and both trend & mean differences. We then spiked in differential trend, mean, or both trend & mean signals into these gold standard genes based on their differential type. To generate the spike-in signals, we selected highly variable genes from the remaining 80% non-gold-standard (non-gs) genes using cells in sample group 0 and using their original unpermuted data. To select highly variable genes, we applied B-splines to fit the relationship between the standard deviation (SD) and the mean of gene expression of the non-gold-standard (non-gs) genes across cells in group 0.

Genes with positive residuals (i.e. SD is larger than its expected value estimated from the mean expression) are selected as highly variable. We applied *k*-means clustering to cluster these genes into 5 clusters using their standardized log2-transformed SAVER-imputed expression. Here the cluster number 5 was determined using the same elbow method as described in TSCAN. For each gene that was clustered, we fit a B-spline on the log2-transformed SAVER-imputed expression against pseudotime. We evaluated the magnitude of change of the gene along pseudotime by calculating a *F* – statistic that compares a full model (which assumes gene expression along pseudotime is modeled using the B-spline curve plus additive noise) and a null model (which assumes gene expression along pseudotime is a constant plus additive noise). We used highly variable genes (i.e. those with positive residuals) as “source genes”. We ordered source genes in increasing *F* – statistics. We categorized the tail 1814 source genes into 4 groups from the smallest to the largest *F* – statistics to represent signal strengths from weakest (1) to highest (4). In each signal-strength simulation, we added the gene expression profiles in each sample from the source genes in the same strength group onto those gold standard genes. The signal-spike-in procedures were performed in SAVER-imputed gene expression matrix and original count matrix in parallel. For gold standard genes with trend difference, we added signals to both group 0 and 1, except that the signals were permuted before adding to group 1. For gold standard genes with mean shift, we permuted the source gene expression profiles within each sample before adding signals to group 0. For gold standard genes with both trend and mean differences, we added source signals directly to group 0 cells without centering the data.

#### 4.11.2 TDE, TCD, and XCD detection

Simulations for evaluating TDE, TCD and XCD detection are presented in Supplementary Notes.

## Declarations

### Acknowledgement

We would like to thank the Maryland Advanced Research Computing Center (MARCC) and The Joint High Performance Computing Exchange (JHPCE) for providing computing resources.

### Availability of data and materials

The data used in this analysis are all publicly available. All data are described in Method Section. All competing methods are described in Additional file 2: Table S1. All code to reproduce the presented analyses are available at https://github.com/Winnie09/trajectory_variability. The `Lamian` package with detailed user manual is publicly available at https://github.com/Winnie09/Lamian. The R package `ggplot2`(v.3.3.0)^41^ for data visualization was used.

### Author contributions

WH, ZJ, SCH and HJ conceived the study. HJ, SCH and WH conceptualized the Lamian framework. WH and HJ developed the statistical model and algorithm with the feedback from ZJ and SCH. WH implemented the model and software. ZJ, ZC and WH prepared the data. WH, ZJ and ZC analyzed the data. WH, ZC, EJW, SCH and HJ interpreted the results. WH, SCH and HJ drafted the manuscript. All authors edited and approved the final manuscript.

### Ethics approval and consent to participate

Not applicable.

### Consent for publication

Not applicable.

### Open Access

This article is distributed under the terms of the MIT License, which is a permissive license with conditions only requiring preservation of copyright and license notices.

## Supplementary materials

Supplementary materials are available as supplementary materials.

**Supplementary file 1 — Supplementary Figures.**

Supplementary Figures S1-S6 (PDF 6.7 MB).

**Supplementary file 2 — Supplementary Notes.**

Supplementary Notes of methods (PDF 128 KB).

**Supplementary file 3 — Supplementary Table S1.**

Comparison of our method Lamian with other pseudotime analyses methods (CSV 1 KB).

