## Supplementary Figures for "A statistical framework for differential pseudotime analysis with multiple single-cell RNA-seq samples"

### Supplementary file 1

### Supplementary Figures

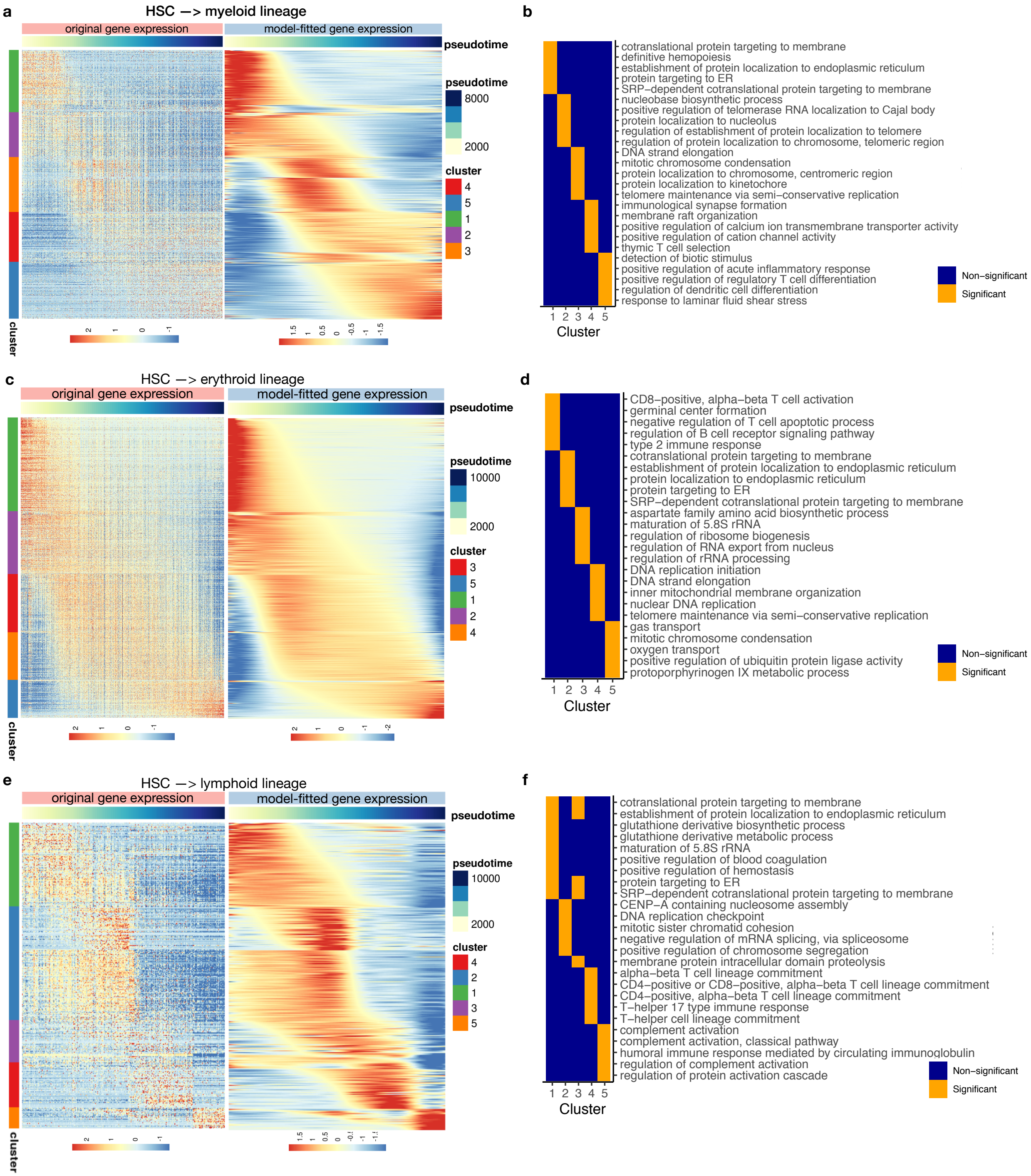


**Figure S1.** Differentially expressed genes along pseudotime found using Lamian's TDE test (Module 3) along the (a,b) myeloid lineage, (c,d) erythroid lineage, and (e,f) lymphoid lineage in the HCA bone marrow scRNA-seq data. (a, c, e) Heatmaps of original gene expression values (left) and model-fitted values (right) along pseudotime. The model-fitted heatmaps are also shown in Fig. 3 (a-c). Rows are TDE genes identified by Lamian using a FDR $< 0.05$ threshold and clustered using $k$-means clustering. Columns are cells ordered by pseudotime. (b, d, f) Heatmaps of enriched gene ontology (GO) terms identified using `topGO`(v.2.36.0)[1] package with $FDR = 0.05$ and $FC > 2$ cutoff for the gene clusters shown in (a, c, e), respectively.

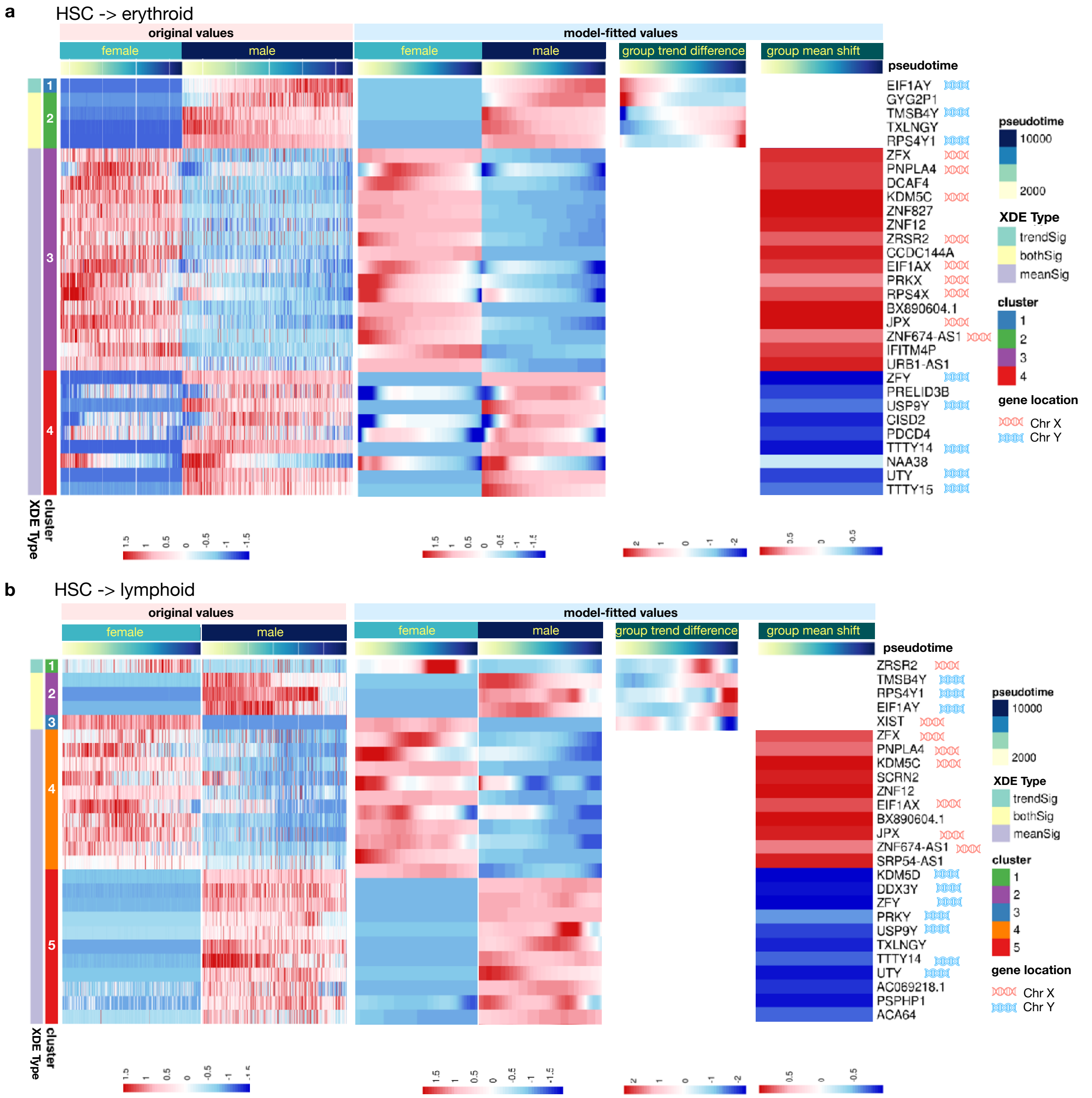


**Figure S2.** Heatmaps of XDE genes along the (a) erythroid and (b) lymphoid lineages associated with sex (male, female) as a covariate found by Lamian's XDE test (Module 3) in the HCA bone marrow scRNA-seq data. Rows are genes identified using FDR < 0.05. Only XDE genes with significant mean shift, trend difference, or both are shown. In the erythroid lineage, 53% (or $n$=17) of the XDE genes identified by Lamian are on sex chromosomes. In the lymphoid lineage, 62% (or $n$=18) of the XDE genes identified by Lamian are on sex chromosomes.

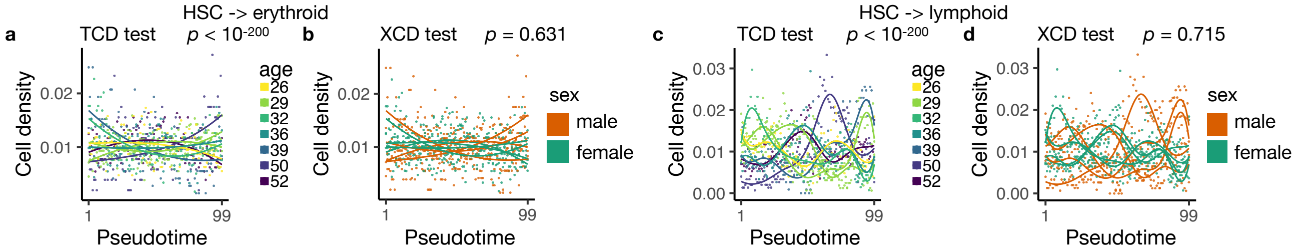


**Figure S3.** TCD and XCD test (Module 4) results for the HCA bone marrow dataset on (a,b) erythroid and (c,d) lymphoid lineages. The pseudotime was split up into 100 equally-spaced bins. Each dot represents the cell density in each bin of one sample (i.e., the number of cells in that bin divided by the total number of cells in that sample). Each curve represents the model-fitted temporal pattern of a sample. In TCD test, the curves are colored by individual samples. In XCD test, they are colored by the sample group.

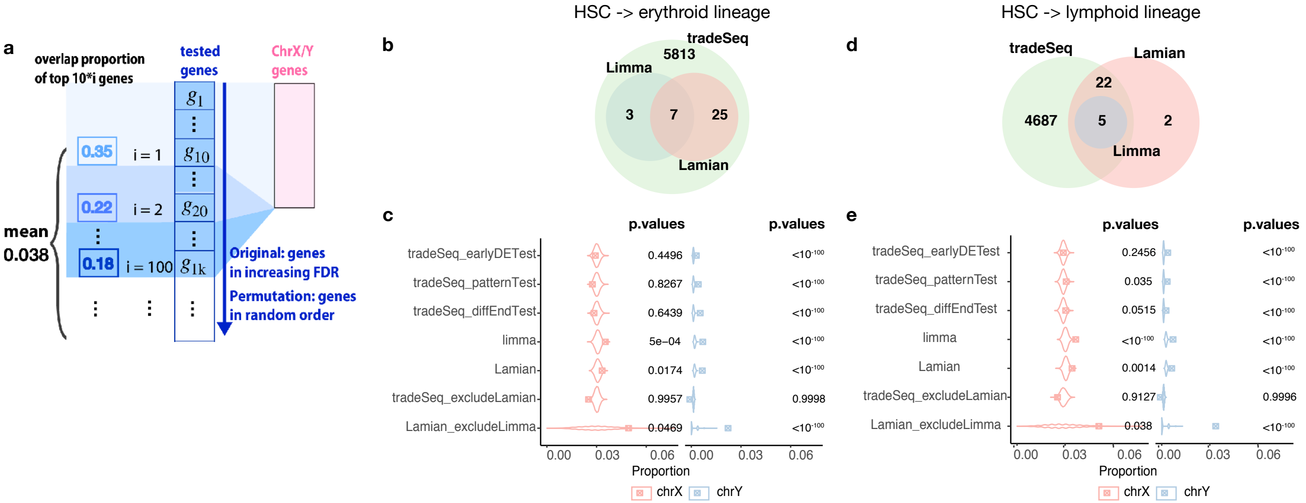


**Figure S4.** Comparison of sex-associated XDE genes reported by different methods in the HCA bone marrow data. (a) Schematic view of the calculation of overlap proportion with sex chromosomes as a evaluation metric in Fig. 5. (b-e) Comparison of XDE genes detected by Lamian and other methods in the erythroid lineage (b-c) and lymphoid lineage (d-e), similar to Fig. 5a,d.

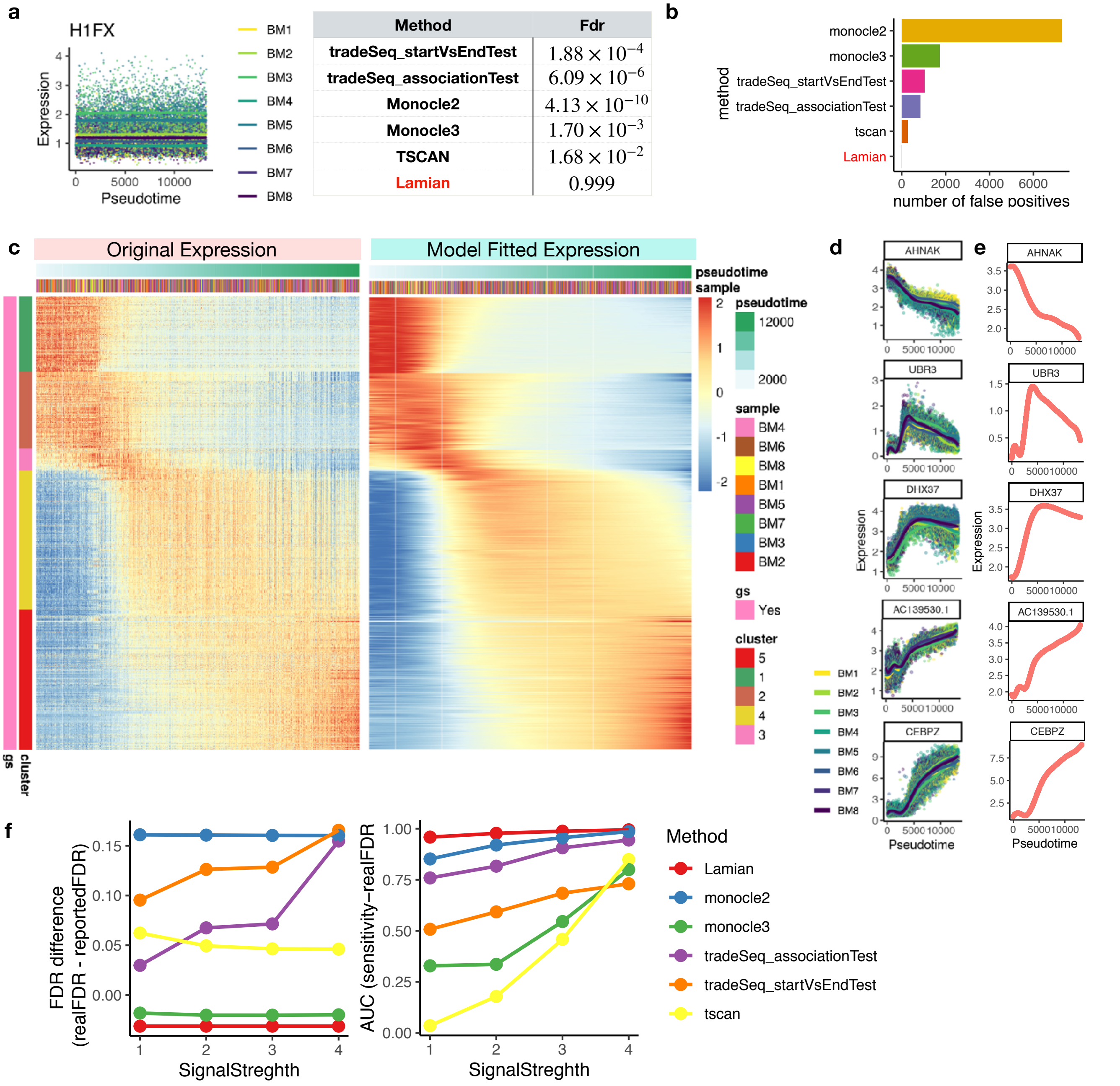


| Method | Fdr |
|---|---|
| tradeSeq_startVsEndTest | $1.88 \times 10^{-4}$ |
| tradeSeq_associationTest | $6.09 \times 10^{-6}$ |
| Monocle2 | $4.13 \times 10^{-10}$ |
| Monocle3 | $1.70 \times 10^{-3}$ |
| TSCAN | $1.68 \times 10^{-2}$ |
| Lamian | 0.999 |

**Figure S5.** Simulation study to assess performance of TDE test. (a-b) illustrate the null simulation. (a) An example null gene without differential expression along pseudotime. Each dot is a cell colored by the sample it belongs to. The gene's expression ($y$-axis) in each cell is shown along pseudotime ($x$-axis). Curves are the model-fitted values for each sample estimated by Lamian. The table shows the FDR reported by Lamian and existing methods about this example gene. (b) Number of false positives reported by each method in the null simulation. (c-f) show results from the spike-in simulation. (c) Heatmap showing expression profiles of TDE genes (rows) along pseudotime-ordered cells (columns) in original values (left) and model-fitted values (right). gs = gold standard TDE genes. Lamian grouped TDE genes into 5 clusters. (d) Five example genes for cluster 1 (top) to 5 (bottom) shown in (c). Each dot is a cell showing the SAVER-imputed expression. The curves are fitted values for each sample. Both are colored by the sample it corresponds to. (e) The population-level gene expression estimates along pseudotime for the five genes in (d). (f) Performance evaluation of all methods. The left plot shows the difference between the true and reported FDR by each method at different signal strength levels. A negative value indicates that the reported FDR properly controls the real FDR. A positive value indicates that the reported FDR underestimate the real FDR. The right plot shows the area under the sensitivity-realFDR curve for different methods. The higher the curve, the more powerful a method is.

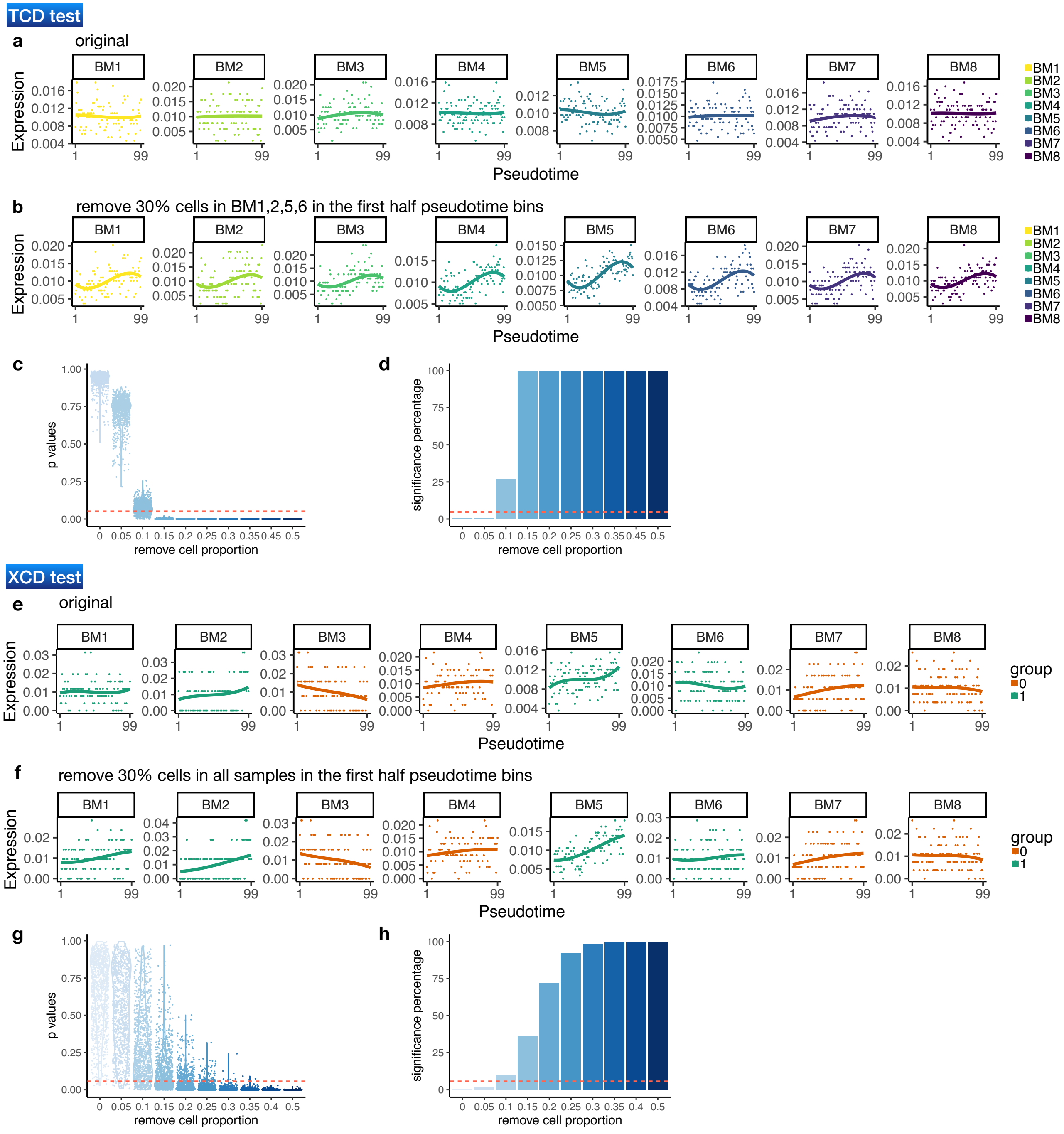


**Figure S6.** Simulation study for evaluating differential cell density tests. (a)-(d) TCD test simulation. (a) Null simulation setting of TCD test. Cell proportion in each of the 100 bins along pseudotime (denoted as dots) and the model-fitted curve for each HCA-BM sample are shown. The pseudotime of the cells have been permuted so that there is no time-dependent pattern. (b) Similar to (a) except that 50% of the cells in the first half pseudotime bins in each of the samples have been removed. (c) A violin plot showing the distribution of $p-$values (dots) obtained from 1,000 simulations where a certain proportion (x-axis) of the cells in the first half pseudotime bins in each sample have been removed. (d) A bar plot showing the percentage of simulations (out of 1,000) that have reported a $p$-value $< 0.05$. (e-h) Similar to (a-d) except that it is in the XCD test simulation setting. Here, instead of removing cells in all samples, we only remove cells in the group 1 samples (BM1,2,5,6); and we do not permute the pseudotime.
