## Supplementary Notes for "A statistical framework for differential pseudotime analysis with multiple single-cell RNA-seq samples"

### 0.1 EM algorithm for fitting the Lamian model

This section presents the Expectation-Maximization(EM) algorithm used to fit the Lamian model for a given gene to estimate the unknown parameters $\Theta = \{\beta, \Omega, \alpha, \eta\}$ and infer $\sigma_s^2$ based on the gene's observed data. Here $\sigma_s^2, \alpha, \eta \in \mathbb{R}$, $\Omega \in \mathbb{R}^{(K+1)\times(K+1)}$, $\beta \in \mathbb{R}^{(K+1)(V+1)}$. In the EM algorithm, we treat sample-level random effects $\mathbf{u} \equiv \{\mathbf{u}_1, \ldots, \mathbf{u}_S\}$ and their variance $\sigma^2 \equiv \{\sigma_1^2, \ldots, \sigma_S^2\}$ as missing data. The observed data are $\mathbf{y} \equiv \{\mathbf{y}_1, \ldots, \mathbf{y}_S\}$. The complete data are $\{\mathbf{y}, \mathbf{u}, \sigma^2\}$.

For a given gene, the complete data likelihood is

$$
\begin{aligned}
L(\Theta) =& P(\mathbf{y}, \mathbf{u}, \sigma^2 | \Theta) \\
=& \prod_{s=1}^{S} \left[ P(\mathbf{y}_s | \mathbf{u}_s, \beta, \sigma_s^2) P(\mathbf{u}_s | \sigma_s^2, \Omega) P(\sigma_s^2 | \alpha, \eta) \right] \\
=& \prod_{s} \Bigg[ (2\pi\sigma_s^2)^{-C_s/2} \exp\left\{ -\frac{1}{2\sigma_s^2} [\mathbf{y}_s - \Phi_s \cdot (\mathbf{X}_s\beta + \mathbf{u}_s)]^T [\mathbf{y}_s - \Phi_s \cdot (\mathbf{X}_s\beta + \mathbf{u}_s)] \right\} \cdot \\
& (2\pi)^{-(K+1)/2} |\sigma_s^2 \Omega|^{-1/2} \exp\left\{ -\frac{1}{2} \mathbf{u}_s^T (\sigma_s^2 \Omega)^{-1} \mathbf{u}_s \right\} \frac{\eta^\alpha}{\Gamma(\alpha)} (\sigma_s^2)^{-\alpha-1} \exp(-\frac{\eta}{\sigma_s^2}) \Bigg] \\
=& \prod_{s} \Bigg[ (2\pi)^{-(C_s+K+1)/2} (\sigma_s^2)^{-(C_s+K+1)/2-\alpha-1} |\Omega|^{-1/2} \frac{\eta^\alpha}{\Gamma(\alpha)} \cdot \\
& \exp\left\{ -\frac{1}{2\sigma_s^2} [(\mathbf{y}_s - \Phi_s \mathbf{X}_s \beta)^T (\mathbf{y}_s - \Phi_s \mathbf{X}_s \beta) - 2(\mathbf{y}_s - \Phi_s \mathbf{X}_s \beta)^T \Phi_s \mathbf{u}_s + \mathbf{u}_s^T (\Phi_s^T \Phi_s + \Omega^{-1}) \mathbf{u}_s + 2\eta] \right\} \Bigg] \\
=& \prod_{s} \left[ (2\pi)^{-(C_s+K+1)/2} (\sigma_s^2)^{-(C_s+K+1)/2-\alpha-1} |\Omega|^{-1/2} \frac{\eta^\alpha}{\Gamma(\alpha)} \cdot \exp\left\{ -\frac{1}{2\sigma_s^2} [\mathbf{L}_s - 2\mathbf{K}_s \mathbf{u}_s + \mathbf{u}_s^T \mathbf{J}_s \mathbf{u}_s + 2\eta] \right\} \right]
\end{aligned}
$$

where

$$
\begin{aligned}
\mathbf{L}_s &= (\mathbf{y}_s - \Phi_s \mathbf{X}_s \beta)^T (\mathbf{y}_s - \Phi_s \mathbf{X}_s \beta) \\
\mathbf{K}_s &= (\mathbf{y}_s - \Phi_s \mathbf{X}_s \beta)^T \Phi_s \\
\mathbf{J}_s &= \Phi_s^T \Phi_s + \Omega^{-1}
\end{aligned}
$$

The complete data log-likelihood is

$$
\begin{aligned}
l(\Theta) = \sum_{s=1}^{S} \Bigg[ & (-\frac{C_s + K + 1}{2} - \alpha - 1) \log(\sigma_s^2) - \frac{1}{2} \log(|\Omega|) + \alpha \log(\eta) - \log(\Gamma(\alpha)) \\
& - \frac{1}{2\sigma_s^2} [\mathbf{L}_s - 2\mathbf{K}_s \mathbf{u}_s + \mathbf{u}_s^T \mathbf{J}_s \mathbf{u}_s + 2\eta] \Bigg] + constant
\end{aligned} \tag{1}
$$

Here the *constant* term does not involve the unknown parameters.

The EM algorithm iterates between the E-step and the M-step below until convergence.

#### 0.1.1 E-step

In iteration $t+1$, the E-step computes the Q-function $Q(\Theta|\Theta^{(t)})$ , which is the expectation of the complete data log-likelihood $l(\Theta)$ with respect to the conditional distribution of missing data $\mathbf{u}$ and $\sigma^2$ given the observed data $\mathbf{y}$ and old parameter values $\Theta^{(t)}$. Based on Equation 1, this involves evaluating (1) $E_{\sigma_s^2|\mathbf{y}_s,\Theta^{(t)}}[\log(\sigma_s^2)]$, (2) $E_{\mathbf{u}_s,\sigma_s^2|\mathbf{y}_s,\Theta^{(t)}}[\frac{\mathbf{u}_s}{\sigma_s^2}]$, and (3) $E_{\mathbf{u}_s,\sigma_s^2|\mathbf{y}_s,\Theta^{(t)}}[\frac{\mathbf{u}_s^T \mathbf{J}_s \mathbf{u}_s}{\sigma_s^2}]$.

Note that

$$
\begin{aligned}
P(\mathbf{u}_s, \sigma_s^2 | \mathbf{y}_s, \Theta) \propto & P(\mathbf{y}_s | \mathbf{u}_s, \sigma_s^2, \Theta) P(\mathbf{u}_s | \sigma_s^2, \Theta) P(\sigma_s^2 | \Theta) \\
\propto & (\sigma_s^2)^{-(C_s+K+1)/2-\alpha-1} \exp\left\{ -\frac{1}{2\sigma_s^2} [\mathbf{L}_s - 2\mathbf{K}_s \mathbf{u}_s + \mathbf{u}_s^T \mathbf{J}_s \mathbf{u}_s + 2\eta] \right\} \\
= & (\sigma_s^2)^{-(C_s+K+1)/2-\alpha-1} \exp\left\{ -\frac{1}{2\sigma_s^2} [\mathbf{L}_s - \mathbf{K}_s \mathbf{J}_s^{-1} \mathbf{K}_s^T + (\mathbf{u}_s - \mathbf{J}_s^{-1} \mathbf{K}_s^T)^T \mathbf{J}_s (\mathbf{u}_s - \mathbf{J}_s^{-1} \mathbf{K}_s^T) + 2\eta] \right\}
\end{aligned}
$$

$$
\begin{aligned}
=&(\sigma_s^2)^{-(C_s+K+1)/2-\alpha-1}\exp\left\{-\frac{1}{2\sigma_s^2}[\mathbf{L}_s-\mathbf{K}_s\mathbf{J}_s^{-1}\mathbf{K}_s^T+2\eta]\right\}\cdot\\
&\exp\left\{-\frac{1}{2\sigma_s^2}[(\mathbf{u}_s-\mathbf{J}_s^{-1}\mathbf{K}_s^T)^T\mathbf{J}_s(\mathbf{u}_s-\mathbf{J}_s^{-1}\mathbf{K}_s^T)]\right\}
\end{aligned}
$$

Thus,

$$
\mathbf{u}_s|\sigma_s^2,\mathbf{y}_s,\Theta\sim N(\mathbf{J}_s^{-1}\mathbf{K}_s^T,\sigma_s^2\mathbf{J}_s^{-1}) \tag{2}
$$

$$
\sigma_s^2|\mathbf{y}_s,\Theta\sim IG(\alpha+C_s/2,\eta+(\mathbf{L}_s-\mathbf{K}_s\mathbf{J}_s^{-1}\mathbf{K}_s^T)/2) \tag{3}
$$

Here the inverse-Gamma distribution for $\sigma_s^2|\mathbf{y}_s,\Theta$ is derived based on

$$
\begin{aligned}
P(\sigma_s^2|\mathbf{y}_s,\Theta)=&\int P(\mathbf{u}_s,\sigma_s^2|\mathbf{y}_s,\Theta)d\mathbf{u}_s\\
&\propto(\sigma_s^2)^{-(C_s+K+1)/2-\alpha-1}\exp\left\{-\frac{1}{2\sigma_s^2}[\mathbf{L}_s-\mathbf{K}_s\mathbf{J}_s^{-1}\mathbf{K}_s^T+2\eta]\right\}|\mathbf{J}_s^{-1}\sigma_s^2|^{1/2}\\
&\propto(\sigma_s^2)^{-C_s/2-\alpha-1}\exp\left\{-\frac{1}{2\sigma_s^2}[\mathbf{L}_s-\mathbf{K}_s\mathbf{J}_s^{-1}\mathbf{K}_s^T+2\eta]\right\}
\end{aligned}
$$

Based on the properties of inverse-Gamma distribution, we have

$$
A_s^{(t)}\equiv E_{\sigma_s^2|\mathbf{y}_s,\Theta^{(t)}}[\log(\sigma_s^2)]=\log(\eta^{(t)}+(\mathbf{L}_s^{(t)}-\mathbf{K}_s^{(t)}(\mathbf{J}_s^{(t)})^{-1}(\mathbf{K}_s^{(t)})^T)/2)-\psi(\alpha^{(t)}+C_s/2) \tag{4}
$$

where $\psi(\cdot)$ is digamma function.

Let

$$
N_s^{(t)}\equiv E_{\sigma_s^2|\mathbf{y}_s,\Theta^{(t)}}[\frac{1}{\sigma_s^2}]=\frac{2\alpha^{(t)}+C_s}{2\eta^{(t)}+\mathbf{L}_s^{(t)}-\mathbf{K}_s^{(t)}(\mathbf{J}_s^{(t)})^{-1}(\mathbf{K}_s^{(t)})^T} \tag{5}
$$

Since $\mathbf{u}_s|\sigma_s^2,\mathbf{y}_s,\Theta\sim N(\mathbf{J}_s^{-1}\mathbf{K}_s^T,\sigma_s^2\mathbf{J}_s^{-1})$, we have

$$
\begin{aligned}
E_{\mathbf{u}_s,\sigma_s^2|\mathbf{y}_s,\Theta^{(t)}}[\frac{\mathbf{u}_s}{\sigma_s^2}]=&E_{\sigma_s^2|\mathbf{y}_s,\Theta^{(t)}}\left[E_{\mathbf{u}_s|\sigma_s^2,\mathbf{y}_s,\Theta^{(t)}}[\frac{\mathbf{u}_s}{\sigma_s^2}]\right]\\
=&E_{\sigma_s^2|\mathbf{y}_s,\Theta^{(t)}}\left[\frac{(\mathbf{J}_s^{(t)})^{-1}(\mathbf{K}_s^{(t)})^T}{\sigma_s^2}\right]\\
=&N_s^{(t)}(\mathbf{J}_s^{(t)})^{-1}(\mathbf{K}_s^{(t)})^T
\end{aligned} \tag{6}
$$

and

$$
\begin{aligned}
E_{\mathbf{u}_s,\sigma_s^2|\mathbf{y}_s,\Theta^{(t)}}[\frac{\mathbf{u}_s^T\mathbf{J}_s\mathbf{u}_s}{\sigma_s^2}]=&E_{\sigma_s^2|\mathbf{y}_s,\Theta^{(t)}}\left[E_{\mathbf{u}_s|\sigma_s^2,\mathbf{y}_s,\Theta^{(t)}}[\frac{\mathbf{u}_s^T\mathbf{J}_s\mathbf{u}_s}{\sigma_s^2}]\right]\\
=&E_{\sigma_s^2|\mathbf{y}_s,\Theta^{(t)}}\left[\frac{1}{\sigma_s^2}\mathbf{K}_s^{(t)}(\mathbf{J}_s^{(t)})^{-1}\mathbf{J}_s(\mathbf{J}_s^{(t)})^{-1}(\mathbf{K}_s^{(t)})^T\right]+tr(\mathbf{J}_s(\mathbf{J}_s^{(t)})^{-1})\\
=&\mathbf{K}_s^{(t)}(\mathbf{J}_s^{(t)})^{-1}\mathbf{J}_s(\mathbf{J}_s^{(t)})^{-1}(\mathbf{K}_s^{(t)})^TE_{\sigma_s^2|\mathbf{y}_s,\Theta^{(t)}}[\frac{1}{\sigma_s^2}]+tr(\mathbf{J}_s(\mathbf{J}_s^{(t)})^{-1})\\
=&N_s^{(t)}\mathbf{K}_s^{(t)}(\mathbf{J}_s^{(t)})^{-1}\mathbf{J}_s(\mathbf{J}_s^{(t)})^{-1}(\mathbf{K}_s^{(t)})^T+tr(\mathbf{J}_s(\mathbf{J}_s^{(t)})^{-1})
\end{aligned} \tag{7}
$$

Based on Equations 4-7, the Q-function is

$$
\begin{aligned}
Q(\Theta|\Theta^{(t)})=&E_{\mathbf{u},\sigma^2|\mathbf{y},\Theta^{(t)}}l(\Theta)\\
=&\sum_s\left[-\alpha A_s^{(t)}-\frac{1}{2}\log(|\Omega|)+\alpha\log(\eta)-\log(\Gamma(\alpha))-\right.
\end{aligned}
$$

$$
\begin{aligned}
&E_{\mathbf{u}_s,\sigma_s^2|\mathbf{y}_s,\Theta^{(t)}}\left(\frac{1}{2\sigma_s^2}[\mathbf{L}_s-2\mathbf{K}_s\mathbf{u}_s+\mathbf{u}_s^T\mathbf{J}_s\mathbf{u}_s+2\eta]\right)\Bigg]+constant\\
=&\sum_s\Bigg[-\alpha A_s^{(t)}-\frac{1}{2}\log(|\Omega|)+\alpha\log(\eta)-\log(\Gamma(\alpha))-\frac{1}{2}N_s^{(t)}\mathbf{L}_s+\mathbf{K}_sN_s^{(t)}(\mathbf{J}_s^{(t)})^{-1}(\mathbf{K}_s^{(t)})^T\\
&-\frac{1}{2}N_s^{(t)}\mathbf{K}_s^{(t)}(\mathbf{J}_s^{(t)})^{-1}\mathbf{J}_s(\mathbf{J}_s^{(t)})^{-1}(\mathbf{K}_s^{(t)})^T-\frac{1}{2}tr(\mathbf{J}_s(\mathbf{J}_s^{(t)})^{-1})-N_s^{(t)}\eta\Bigg]+constant\\
=&\sum_s\Bigg[-\alpha A_s^{(t)}-\frac{1}{2}\log(|\Omega|)+\alpha\log(\eta)-\log(\Gamma(\alpha))\\
&+N_s^{(t)}\left(-\frac{1}{2}(\Phi_s\mathbf{X}_s\beta)^T(\Phi_s\mathbf{X}_s\beta)+(\Phi_s\mathbf{X}_s\beta)^T\mathbf{y}_s-(\Phi_s\mathbf{X}_s\beta)^T\Phi_s(\mathbf{J}_s^{(t)})^{-1}(\mathbf{K}_s^{(t)})^T-\eta\right)\\
&-\frac{1}{2}\left(tr(\Omega^{-1}(\mathbf{J}_s^{(t)})^{-1})+N_s^{(t)}\mathbf{K}_s^{(t)}(\mathbf{J}_s^{(t)})^{-1}\Omega^{-1}(\mathbf{J}_s^{(t)})^{-1}(\mathbf{K}_s^{(t)})^T\right)\Bigg]+constant
\end{aligned}
\tag{8}
$$

##### *0.1.2 M-step*

By maximizing the Q-function with respect to the unknown parameters $\Theta$, we obtain the new parameter estimates.

- $\eta^{(t+1)}$: it is the solution to

$$\log\eta=\sum_s A_s^{(t)}/S+\psi(\eta\sum_s N_s^{(t)}/S) \tag{9}$$

  where $\psi(\cdot)$ is the digamma function. This can be solved using bound constrained optimization[1].

- $\alpha^{(t+1)}$:

$$\alpha^{(t+1)}=\eta^{(t+1)}\sum_s N_s^{(t)}/S \tag{10}$$

- $\beta^{(t+1)}$:

$$\beta^{(t+1)}=\left(\sum_s N_s^{(t)}(\Phi_s\mathbf{X}_s)^T(\Phi_s\mathbf{X}_s)\right)^{-1}\left(\sum_s N_s^{(t)}\left[(\Phi_s\mathbf{X}_s)^T(\mathbf{y}_s-\Phi_s(\mathbf{J}_s^{(t)})^{-1}(\mathbf{K}_s^{(t)})^T)\right]\right) \tag{11}$$

- $\Omega^{(t+1)}$:

$$\Omega^{(t+1)}=\sum_s\left[(\mathbf{J}_s^{(t)})^{-1}+N_s^{(t)}(\mathbf{J}_s^{(t)})^{-1}(\mathbf{K}_s^{(t)})^T(\mathbf{K}_s^{(t)})(\mathbf{J}_s^{(t)})^{-1}\right]/S \tag{12}$$

#### 0.2 Observed data likelihood

To compare two models, the likelihood ratio statistic is computed based on the observed data likelihood. The observed data likelihood for a Lamian model is:

$$L_{obs}(\Theta)=P(\mathbf{y}|\Theta)=\prod_s P(\mathbf{y}_s|\Theta) \tag{13}$$

Here

$$
\begin{aligned}
P(\mathbf{y}_s|\Theta)=&\int\int P(\mathbf{y}_s,\mathbf{u}_s,\sigma_s^2|\Theta)\,\mathrm{d}\mathbf{u}_s\,\mathrm{d}\sigma_s^2\\
=&\int\int P(\mathbf{y}_s|\mathbf{u}_s,\beta,\sigma_s^2)P(\mathbf{u}_s|\sigma_s^2,\Omega)P(\sigma_s^2|\alpha,\eta)\,\mathrm{d}\mathbf{u}_s\,\mathrm{d}\sigma_s^2\\
=&\int\int\Bigg[(2\pi)^{-(C_s+K+1)/2}(\sigma_s^2)^{-(C_s+K+1)/2-\alpha-1}|\Omega|^{-1/2}\frac{(\eta)^\alpha}{\Gamma(\alpha)}\\
&\exp\left\{-\frac{1}{2\sigma_s^2}[\mathbf{L}_s-2\mathbf{K}_s\mathbf{u}_s+\mathbf{u}_s^T\mathbf{J}_s\mathbf{u}_s+2\eta]\right\}\Bigg]\,\mathrm{d}\sigma_s^2\,\mathrm{d}\mathbf{u}_s
\end{aligned}
$$

$$
\begin{aligned}
=&(2\pi)^{-(C_s+K+1)/2}|\Omega|^{-1/2}\frac{(\eta)^\alpha}{\Gamma(\alpha)}\Gamma\left(\frac{C_s+K+1}{2}+\alpha\right)\\
&\int\left(\frac{1}{2}[\mathbf{L}_s-2\mathbf{K}_s\mathbf{u}_s+\mathbf{u}_s^T\mathbf{J}_s\mathbf{u}_s+2\eta]\right)^{-(C_s+K+1)/2-\alpha}\mathrm{d}\mathbf{u}_s\\
=&(2\pi)^{-(C_s+K+1)/2}|\Omega|^{-1/2}\frac{(\eta)^\alpha}{\Gamma(\alpha)}\Gamma\left(\frac{C_s+K+1}{2}+\alpha\right)\\
&\int\left(\frac{1}{2}[\mathbf{L}_s+2\eta+(\mathbf{u}_s-\mathbf{J}_s^{-1}\mathbf{K}_s^T)^T\mathbf{J}_s(\mathbf{u}_s-\mathbf{J}_s^{-1}\mathbf{K}_s^T)-\mathbf{K}_s\mathbf{J}_s^{-1}\mathbf{K}_s^T]\right)^{-(C_s+K+1)/2-\alpha}\mathrm{d}\mathbf{u}_s\\
=&(2\pi)^{-(C_s+K+1)/2}|\Omega|^{-1/2}\frac{(\eta)^\alpha}{\Gamma(\alpha)}\Gamma\left(\frac{C_s+K+1}{2}+\alpha\right)2^{(C_s+K+1)/2+\alpha}\\
&\int\left(\mathbf{L}_s+2\eta-\mathbf{K}_s\mathbf{J}_s^{-1}\mathbf{K}_s^T+(\mathbf{u}_s-\mathbf{J}_s^{-1}\mathbf{K}_s^T)^T\mathbf{J}_s(\mathbf{u}_s-\mathbf{J}_s^{-1}\mathbf{K}_s^T)\right)^{-(C_s+K+1)/2-\alpha}\mathrm{d}\mathbf{u}_s\\
&\text{(Note the connection between the integrand and the probability density function of multivariate t-distribution)}\\
=&(2\pi)^{-(C_s+K+1)/2}|\Omega|^{-1/2}\frac{(\eta)^\alpha}{\Gamma(\alpha)}\Gamma\left(\frac{C_s+K+1}{2}+\alpha\right)2^{(C_s+K+1)/2+\alpha}\\
&(\mathbf{L}_s+2\eta-\mathbf{K}_s\mathbf{J}_s^{-1}\mathbf{K}_s^T)^{-(C_s+K+1)/2-\alpha}\frac{\Gamma\left(\frac{C_s+2\alpha}{2}\right)(C_s+2\alpha)^{(K+1)/2}\pi^{(K+1)/2}}{\Gamma\left(\frac{C_s+K+1}{2}+\alpha\right)}\\
&\left|\frac{(\mathbf{L}_s+2\eta-\mathbf{K}_s\mathbf{J}_s^{-1}\mathbf{K}_s^T)(\mathbf{J}_s)^{-1}}{C_S+2\alpha}\right|^{1/2}\\
=&\frac{(2\eta)^\alpha\Gamma(C_s/2+\alpha)}{(\pi)^{C_s/2}\Gamma(\alpha)|\Omega|^{1/2}|\mathbf{J}_s|^{1/2}(\mathbf{L}_s+2\eta-\mathbf{K}_s\mathbf{J}_s^{-1}\mathbf{K}_s^T)^{C_s/2+\alpha}}
\end{aligned}
$$

Thus, the log of the observed data likelihood used to construct the likelihood ratio test statistic is

$$
\begin{aligned}
l_{obs}(\Theta)=&\sum_s\log P(\mathbf{y}_s|\Theta)\\
=&\sum_s\left[\alpha\log(2\eta)+\log\Gamma\left(\frac{C_s}{2}+\alpha\right)-\frac{C_s}{2}\log(\pi)-\log\Gamma(\alpha)\right.\\
&\left.-\frac{1}{2}\log(|\Omega|)-\frac{1}{2}\log(|\mathbf{J}_s|)-\left(\frac{C_s}{2}+\alpha\right)\log(\mathbf{L}_s+2\eta-\mathbf{K}_s\mathbf{J}_s^{-1}\mathbf{K}_s^T)\right]
\end{aligned}
\tag{14}
$$

where the parameters $\Theta$ are set to be the maximum likelihood estimates (MLE) obtained from the EM algorithm (i.e., $\Theta^{(t)}$ from the last EM iteration).

### 0.3 Evaluation of TDE detection

We compared Lamian with several state-of-the-art TDE detection methods including Monocle2/3, tradeSeq (which is the TDE method used by Slingshot), and TSCAN.

We first created a null simulation (TDE simulation 1) using the bone marrow data by permuting cells' pseudotime within each sample. This creates a dataset where no genes are differential along pseudotime but sample-level variation is retained (Fig. S5a). Instead of considering the variability across samples, existing methods analyze all cells as if they were from a single sample. Therefore, they all reported a large number of false positives at the claimed 5% FDR cutoff. By contrast, Lamian successfully controlled the FDR and did not report any false positive (Fig. S5a,b).

In another analysis (TDE simulation 2) which builds upon the null simulation above, we added spike-in signals *in silico* with varying signal-to-noise ratio to a random set of genes which provide gold standard TDE genes. In other words, after creating the null simulation data above which contain no TDE genes, we randomly selected 20% (1814) genes to spike in signals to create the gold standard TDE genes (gs genes). To create the spike-in signals, we first selected source genes in the same way as in the XDE simulation. Next, the source genes were categorized into four groups from the weakest signal strength (group 1) to the highest signal strength (group 4), using a similar procedure to that in the spike-in simulation in XDE detection, except that all cells were used here instead of only cells in sample group 0. In each sample, we added the gene expression profiles of the source genes in the same strength group to those of the gold standard TDE genes. The signal spike-in step was operated in SAVER-imputed values and counts in parallel. This results in a dataset where we know which genes are TDE. At the same time, the sample-level variability in real data is also retained. The spike-in signals were simulated from 5 different patterns, which were successfully recovered by Lamian via unsupervised clustering of TDE genes (Fig. S5c-e). Compared with other TDE detection methods, Lamian not only offered the highest sensitivity to detect TDE genes but also controlled FDR (i.e. True FDR - Reported FDR $< 0$). By contrast, the other methods either failed to control FDR or had lower power as measured by the area under the sensitivity vs. true FDR curve (AUC) (Fig. S5f).

#### 0.4 Evaluation of TCD detection

To evaluate TCD detection, we first created a null dataset by randomly permuting the pseudotime time of the cells in erythroid lineage within each sample of the HCA bone marrow dataset. After permutation, there is no temporal variation expected (Fig. S6a). Next, we divided the pseudotime into 100 non-overlapping bins. To add spike-in signals, we randomly excluded $x\%$ ($x = 0, 5, 10, \ldots, 50$) cells from each of the samples in the first half of the pseudotime bins. This results in cell density changes along pseudotime where the cell density in the first half of the bins is expected to drop (Fig. S6b). Increasing $x$ results in a larger drop. For each $x$ value, we repeated the simulation 1000 times and applied TCD test to each simulation dataset, resulting in 1000 $p$-values. The distribution of the $p$-values for each $x$ is shown in Fig. S6c. The p-values became smaller when excluding more cells (larger $x$) (Fig. S6c). Fig. S6d shows the percentage of p-values that were smaller than the $\alpha$=0.05 significance cutoff. When no cells were excluded ($x = 0$), one does not expect any cell density change along pseudotime. Indeed, we observed that less than 5% p-values from TCD tests were below 0.05, indicating that TCD test correctly controlled the Type I error rate. When cells were excluded with increasing proportion (i.e., increasing $x$), the percentage of p-values that were below 0.05 also increased, indicating an increasing power of the TCD test for detecting increasingly larger cell density changes along pseudotime (Fig. S6d).

#### 0.5 Evaluation of XCD detection

To evaluate XCD detection, we begin with creating a null dataset. To so do, the eight HCA bone marrow samples were randomly partitioned into two groups (BM1,2,5,6 as group 1, and the remaining samples as group 0). We divided the pseudotime in erythroid lineage into 100 non-overlapping bins. Within each bin and for each sample, we calculated the proportion of cells falling into that bin (Fig. S6e). For each bin, the median of cell proportions within each sample group was calculated. For the sample group with a larger median cell proportion, we randomly excluded cells from that group so that the two sample groups had the same median cell proportion after exclusion. This results in a null dataset without cell density differences between the two sample groups (Fig. S6e). We then randomly bootstrapped the cells and created 1000 null datasets.

To add spike-in differential signals to null data, we randomly excluded $x\%$ ($x = 0, 5, 10, \ldots, 50$) cells from each sample in one sample group and in the first half of the pseudotime bins. We repeated the process of excluding cells 1000 times, resulting in 1000 simulation datasets for each $x$ (Fig. S6f). We applied XCD test to each dataset. The distribution of the $p$-values for each $x$ is shown in Fig. S6g. The p-values became smaller when the group difference becomes larger (i.e. when $x$ increases). Fig. S6h shows the percentage of p-values that were smaller than the $\alpha$=0.05 significance cutoff. When there is no group difference ($x = 0$), less than 5% p-values from XCD tests were below 0.05, indicating that XCD test correctly controlled the Type I error rate. With increasing group difference (i.e., increasing $x$), the percentage of p-values that were below 0.05 also increased, indicating an increasing power of the XCD test (Fig. S6h).
